# Guidelines for cell-type heterogeneity quantification based on a comparative analysis of reference-free DNA methylation deconvolution software

**DOI:** 10.1101/698050

**Authors:** Clementine Decamps, Florian Privé, Raphael Bacher, Daniel Jost, Arthur Waguet, HADACA consortium, Eugene Andres Houseman, Eugene Lurie, Pavlo Lutsik, Aleksandar Milosavljevic, Michael Scherer, Michael G.B. Blum, Magali Richard

**Affiliations:** Laboratory TIMC-IMAG, UMR 5525, Univ. Grenoble Alpes, CNRS, F-38700 Grenoble, France; Independent Statistical Consultant, La Center, WA, USA; Bioinformatics Research Laboratory, Molecular and Human Genetics Department, Baylor College of Medicine, Houston, Texas, USA; Division of Cancer Epigenomics, German Cancer Research Center (DKFZ), Heidelberg, Germany; Department of Genetics/Epigenetics, Saarland University, 66123 Saarbruecken, Germany

## Abstract

Cell-type heterogeneity of tumors is a key factor in tumor progression and response to chemotherapy. Tumor cell-type heterogeneity, defined as the proportion of the various cell-types in a tumor, can be inferred from DNA methylation of surgical specimens. However, confounding factors known to associate with methylation values, such as age and sex, complicate accurate inference of cell-type proportions. While reference-free algorithms have been developed to infer cell-type proportions from DNA methylation, a comparative evaluation of the performance of these methods is still lacking.

Here we use simulations to evaluate several computational pipelines based on the software packages MeDeCom, EDec, and RefFreeEWAS. We identify that accounting for confounders, feature selection, and the choice of the number of estimated cell types are critical steps for inferring cell-type proportions. We find that removal of methylation probes which are correlated with confounder variables reduces the error of inference by 30-35%, and that selection of cell-type informative probes has similar effect. We show that Cattell’s rule based on the scree plot is a powerful tool to determine the number of cell-types. Once the pre-treatment steps are achieved, the three deconvolution methods provide comparable results. We observe that all the algorithms’ performance improves when inter-sample variation of cell-type proportions is large or when the number of available samples is large. We find that under specific circumstances the methods are sensitive to the initialization method, suggesting that averaging different solutions or optimizing initialization is an avenue for future research. Based on the lessons learned, to facilitate pipeline validation and catalyze further pipeline improvement by the community, we develop a benchmark pipeline for inference of cell-type proportions and implement it in the R package *medepir*.

## Background

Since the development of high-throughput sequencing technologies, cancer research has focused on characterizing genetic and epigenetic changes that contribute to the disease. However, these studies often neglect the fact that tumors are constituted of cells with different identities and origins (cell heterogeneity) (Alizadeh et al. 2015). Quantification of tumor heterogeneity is of utmost interest as multiple components of a tumor are key factors in tumor progression and response to chemotherapy (Alizadeh et al. 2015).

Advanced microdissection techniques to isolate a population of interest from heterogeneous clinical tissue samples are still not feasible in daily practice (too complicated and costly). An alternative is to rely on computational deconvolution methods that infer cell-type composition. Recently, several “reference-free” algorithms have been proposed to estimate tumor cell-type heterogeneity from global DNA methylation profiling of surgical specimens (Houseman et al. 2016; Lutsik et al. 2017; Onuchic et al. 2016). Indeed, DNA methylation is a stable molecular marker with a cell type-specific profile dynamically acquired during cell differentiation (Jones 2012; Roadmap Epigenomics Consortium et al. 2015) and thus provides valuable information for cell-type heterogeneity characterization and quantification. The “reference-free” algorithms are labelled as such because they do not require *a priori* information about DNA methylation profiles of cell types found within tumors: they directly infer them from DNA methylation samples using computational methods. While not requiring the *a priori* reference information, some algorithms are designed to use such information when available (Houseman et al. 2016; Lutsik et al. 2017; Onuchic et al. 2016). In the absence of reference information, confounding factors affecting methylation values, such as age and sex, can potentially influence the inference of cell-type proportions. Moreover, the heuristics that are used to estimate the number of underlying cell types may differ between each deconvolution method and the sensitivity of the methods to such variability remains uncharacterized. Therefore, there is an urgent need to comprehensively characterize the current analysis pipelines, identify key features influencing their performance and provide benchmarks and recommendations to guide the application and further development of pipelines that quantify cell-type heterogeneity from reference-free DNA methylation samples (Titus et al. 2017).

Methods correcting for cell-type heterogeneity have already been compared for their statistical power to detect significant associations between epigenetic variation and biological traits (McGregor et al. 2016)(Kaushal et al. 2017). When associating epigenetic variation to phenotypic traits (Epigenome Wide Association Studies, EWAS), cell-type proportions are considered as confounding factors, their inference is not the main objective, but rather an intermediate step that can contribute to reducing false positive associations. In contrast, we here compare reference-free deconvolution methods with the estimation of cell-type proportions as the main objective, as they are directly related to tumorigenesis (Alizadeh et al. 2015). This objective excludes several software packages from our comparison that instead return latent or surrogate variables, which are not interpretable in terms of cell-type proportions (McGregor et al. 2016).

We compare three software packages that infer cell type proportions based on methylation data: MeDeCom, EDec, and RefFreeEWAS (Onuchic et al. 2016; Houseman et al. 2016; Lutsik et al. 2017). For our comparisons, we rely on simulations where real methylation profiles of different cell types are mixed in differing proportions. While some of the methods include series of steps that may be considered a pipeline, the simulations focus on comparing the core deconvolution step shared by all the three methods (e.g., Stage 1 of EDec) that solves a convolution equation that contains two key variables: (i) the cell-type proportions within the samples, and (ii) the average methylation profiles of constituent cell types. The main outcome of this core deconvolution step are estimates of cell-type proportions and of the methylation profiles of constituent cell types, which are needed to characterize the constituent cell types and quantify tumor heterogeneity. Because accurate references for cell-type specific methylation profiles are sparse, especially for solid tissues and cancer cell types, we further assume that reference data for constituent cell-types is not available, which excludes reference-based methods from our comparative analysis (Teschendorff et al. 2017; Zheng et al. 2018).

We here evaluate key factors affecting performance of deconvolution pipelines. We examine to what extent cell-type proportions can be accurately inferred when accounting for measured confounding factors. We determine how feature selection impacts algorithms’ performance at inferring cell-type proportions. We ask how variable are the results of deconvolution to the randomly selected initialization of local optimization involved in solving deconvolution equation. We also test several methods for selecting appropriate number of constituent cell types and ask how sensitive the results are to the variation in cell type number. Based on these, we provide general guidelines for the development of reference-free deconvolution pipelines and define a benchmark pipeline to catalyze further application and improvement of reference-free deconvolution methods.

## Results

### Evaluation of computational frameworks to estimate cell type composition

We apply MeDeCom, EDec (Stage 1, the core deconvolution step), and RefFreeEWAS to estimate heterogeneity within simulated tumorous tissues (Lutsik et al. 2017; Onuchic et al. 2016; Houseman, Molitor, and Marsit 2014). Simulations are encoded in a matrix D of size MxN, where M represents the number of CpG probes and N represents the number of samples. All these software packages perform various types of non-negative matrix factorization to infer cell type proportions (matrix A of size KxN, with K as the putative number of cell types) and cell type-specific methylation profiles (matrix T of size MxK) by solving D = TA, or rather by minimizing, under various constraints (that vary between the three tested algorithms), the error term: ‖ *D* – *TA* ‖_2_(see Material and Methods). We simulate D with 5 cell types (K = 5): 2 cancer-like cells (lung epithelial and mesenchymal), healthy epithelial cells (lung epithelial), immune cells (T lymphocytes), and stromal cells (fibroblasts). These simulations mainly depend on a parameter α0, which controls the diversity of the generated samples (see Material and Methods): When α_0_ is small (∼1), the simulated proportions of the K cell-types are diverse among samples and as α_0_ increases, the variability decreases to the point at which proportions are the same for all samples. Finally, we simulate the effect of confounding factors on these mixtures by using a regression model of methylation data computed from real lung cancer clinical datasets (Supplementary Figure S1, see Material and Methods for details).

To evaluate the methods performance, we use Mean Absolute Error (MAE, see Material and Methods) as a metric to compare inferred individual cell type proportions to the ground truth. First, we tested the effect of altering four simulation parameters on the methods performance (Figure 1 and Supplementary Figure S1): (1) the number of simulated samples (N, ranging from 10 to 500), (2) the inter-sample variation in mixture proportions (α_0_, from 1 to 10,000), (3) the magnitude of random noise added to the mixture component (*ε*, from 0.05 to 0.2) and (4) the set of K cell types used to simulate complex tissues (termed as the cell line background, G, which includes all specificities related to the cell line establishment, such as the genetic background of the donor) (see Material and Methods for details). As expected, increasing the sample size (Supplementary Figure S2) or decreasing random noise improves the performance of all methods (Figure 1). By contrast, their performance is not sensitive to changes of the cell background (Figure 1). Increasing inter-sample proportion variability also substantially improves performance of all methods (Figure 1). Average error (mean error across the three methods) is 0.074 (α_0_ = 1) when inter-sample variation is large, increases to 0.147 (α_0_ = 10) when variation is moderate, and reaches 0.194 (α_0_ = 100) when variation is almost zero (Figure 1 and Supplementary Figure S3). In this first direct comparison, the three deconvolution methods account for all 23,381 probes corresponding to a subset of the Illumina 27k DNA methylation probes, with no specific filtering. To run the algorithms, we used the following functions and parameters: RefFreeEWAS::RefFreeCellMix (5 cell types, 9 iterations), EDec::run_edec_stage_1 (5 cell types, all probes kept as informative loci, maximum iterations = 2000), and MeDeCom::runMeDeCom (5 cell types, lambdas in 0, 0.00001, 0.0001, 0.001, 0.01, 0.1), maximum iterations = 300, 10 random initializations, number of cross-validation folds = 10). Under these not-optimized conditions (i.e. with no pre-processing steps), we observe that all methods provide comparable performance, each algorithm performing best under specific conditions and parameter settings. RefFreeEwas performs best for 9 out of 17 different parameter settings, MeDeCom for 6, and EDec for 2 conditions. Error obtained with EDec is on average 8% larger than the error obtained with RefFreeEwas and 2% larger than MeDeCom.

**Figure 1:**
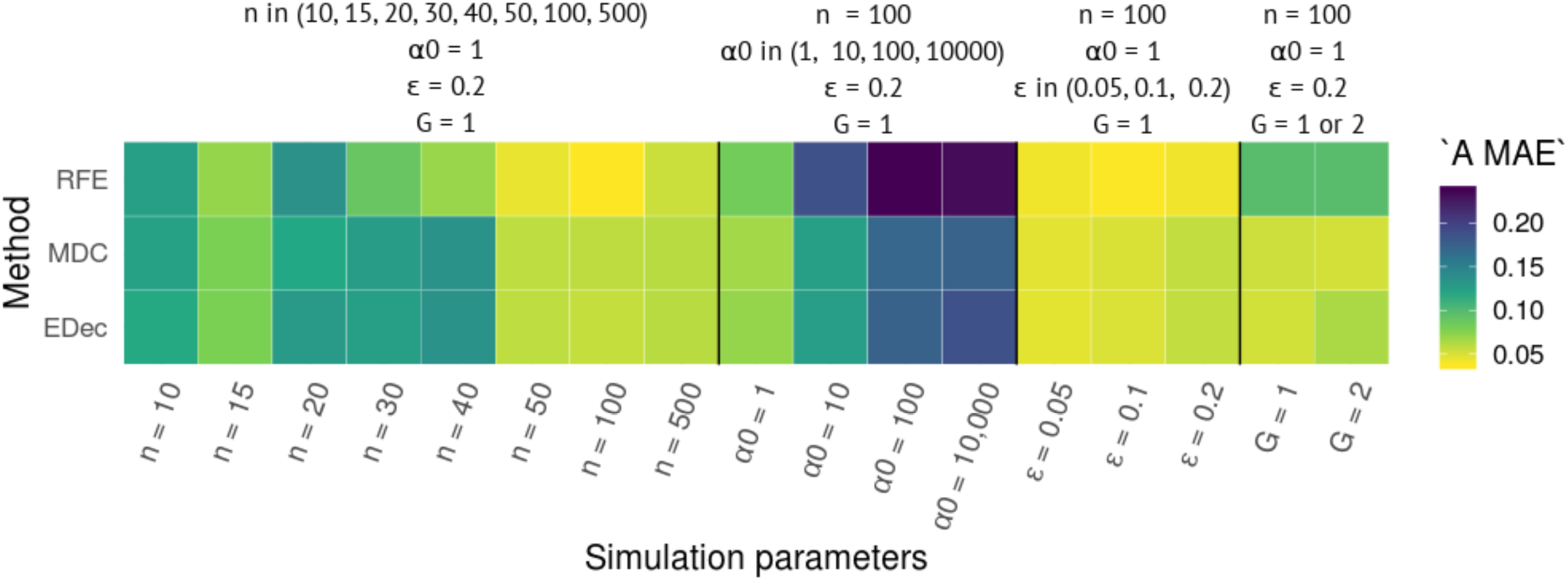
Performance of the 3 deconvolution methods for different parameter settings. Heatmap of method performance (‘A MAE’: Mean Absolute Error on estimated A, the matrix of cell proportions). RFE stands for RefFreeEWAS, MDC for MeDeCom and EDec for EDec stage 1. All algorithms were run on 10 D matrices corresponding to 10 different realizations of the random *ε*-controlled process on one D matrix computed from one simulated A matrix, each time, with the following parameters n (number of samples), α0 (inter-sample variation in mixture proportion), *ε* (magnitude of random noise applied on D) and G (the cell line background of the cell lineages). A random A matrix was used for testing the effect of G1 and G2, another random A matrix was used for testing the effect of *ε* magnitude. Testing the effect of n and α0 required independent simulation of A each time. As a consequence, the four simulations corresponding to the set of parameters n = 100, α_0_ = 1, *ε* = 0.2, G = 1 have different results, because these simulations are based on different randomly simulated A matrices (see figure 7 for a systematic analysis of performance variation according to the random simulations of A).

These results suggest that the differences between the tested algorithms are minor when default parameters are used and no filters are applied on the provided DNA methylation probes. The main variations in performance are related to simulation parameters, such as sample size (n) or inter-sample proportion variability (α0).

### Different strategies to initialize matrix factorization

Optimization algorithms implemented in MeDeCom, EDec and RefFreeEWAS start with an initial condition for either T, the cell type-specific methylation matrix, or A, the cell type proportion matrix. RefFreeEWAS initializes the T matrix. To explore the role of T initialization, we run the RefFreeEWAS package on 10 D matrices (generated from 10 random simulations of an A matrix with the following parameters: K = 5, n = 100, α0 = 1 and *ε* = 0.2), using the following initialization schemes (see Material and Methods for details). K averaged methylation profiles are derived from the D matrix either by hierarchical clustering (estimation of the mean methylation of the K first clusters using a complete linkage method) based on (1) Euclidean or (2) Manhattan distances; either by (3) singular value decomposition (SVD) (corresponding to the K highest singular values) with discretized methylation values (0/1); (4) the ground truth corresponding to real T matrix used in the simulations is also tested (4) (Figure 2, Supplementary table S1). The RefFreeEWAS outcome varies significantly according to how it is initialized, especially when T is initialized by hierarchical clustering applied on the D matrix (estimation of the mean methylation of the K first clusters). Indeed, these two approaches based on hierarchical clustering display a high variability depending on the random simulations of A. Hierarchical clustering based on Euclidean distance or Manhattan distance performs similarly (error ranging from 0.022 to 0.135 for Euclidean distance, and from 0.022 to 0.138 for Manhattan distance). In some cases, they even outcompete initialization with the real T matrix used for simulations (0.059 mean MAE for real T), whereas the SVD approach perform systematically worse (0.152 mean MAE). Surprisingly, the effect of T initialization is highly dependent on the inter-sample variation in mixture proportion (α0). When variation of proportion is low (α0 = 10,000), SVD initialization is more efficient than hierarchical clustering (0.077 mean MAE for SVD-based initialization versus 0.23 mean MAE for clustering-based initialization) (Supplementary Figure S4, Supplementary table S2).

**Figure 2:**
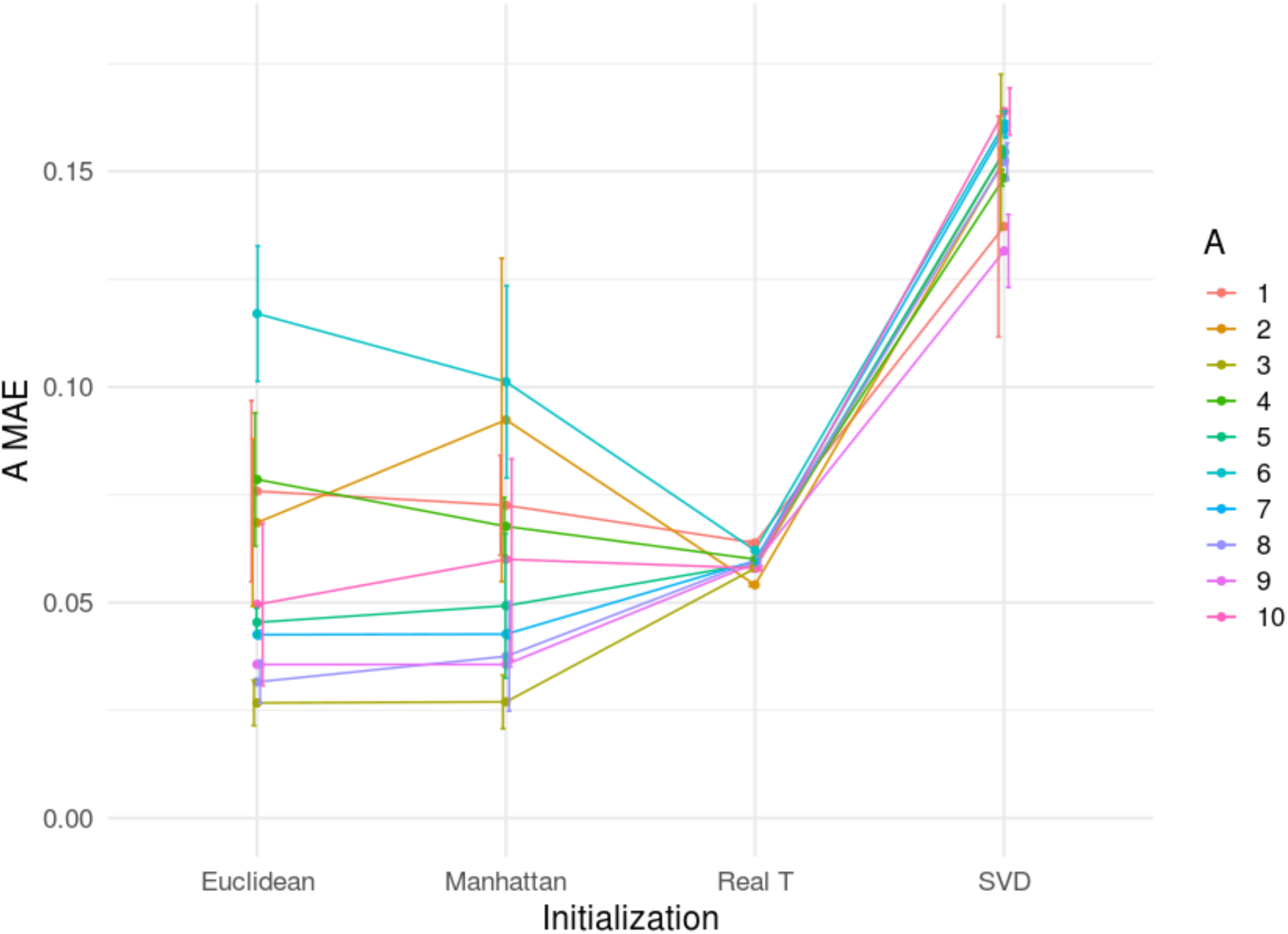
Impact of algorithm initialization of RefFreeEWAS method performance. ‘A MAE’ is shown for 10 D matrices (mean value of 10 random noises applied on D) computed from 10 random A. Each colour represents a different simulated A. Error bars represent standard deviation on 10 random noises. The following parameters were used to simulate D: K = 5, α0 = 1, *ε* = 0.2, G = 1 and n = 100). Euclidean corresponds to RefFreeEWAS::RefFreeCellMixInitialize function applied with the default parameter dist.method = “euclidean”. Manhattan corresponds to RefFreeEWAS::RefFreeCellMixInitialize function applied with the parameter dist.method = “manhattan”. Real T corresponds to RefFreeEWAS::RefFreeCellMix used with the parameter mu0 = real_T, with real_T the matrix composed of the 5 cell types used to simulate D. SVD corresponds to RefFreeEWAS::RefFreeCellMixInitializeBySVD function with default parameters.

MeDeCom performs multiple random initializations of A, the cell type proportion matrix, and EDec performs initialization with random guesses of proportions of cell types in each sample (randomized A generated from a Dirichlet distribution meeting boundary conditions of 0≤A≤1). To roughly estimate the differences between these two approaches on the outcome of the deconvolution algorithms, we run MeDeCom (without the regularization function: parameter lambda = 0) and EDec on 10 independent D matrices (generated from 10 random simulation of matrices A, n = 100, α0 = 1 and *ε* = 0.2) (Supplementary Figure S5). Interestingly, both approaches give similar results. There appears to be a range of errors across 10 different D matrices simulated for both methods tested, with 3 D matrices showing a high standard deviation across 10 random noises (Supplementary Figure S5), suggesting these initialization methods may show sensitivity as well in certain situations.

In summary, the strategies of initialization can have important impacts on method performance, some of them being highly dependent on the composition of the original D matrix. Further work is therefore needed to better understand the relationship between the initialization of the algorithm and method performance.

### Feature selection and accounting for confounders

Variation in DNA methylation is associated with different factors (such as age, sex, batch effects, etc.) that are not always related with cell-type composition. A popular assumption is that removing probes by feature selection will improve performance of deconvolution methods. Yet, such an approach may also discard relevant biological information (Teschendorff et al. 2017).

Because probe selection always involves probe removal, we evaluate to what extent removing probes (i.e. Feature Selection, FS) impacts estimation error. In the previous section, all the 23,381 probes were used by the three non-negative matrix factorization algorithms. Here, we apply different types of feature selection and measure their impacts on deconvolution errors. First, we perform feature selection (FS) without removal of confounding probes (Figure 3A and B, left panel). When keeping only the most variable probes, or the ones that are the most correlated with principal components (PCs), error remains similar to when using no FS (FS variance and FS PCA, resp. in Figure 3). When keeping only cell-type variable informative probes based on literature curation and use of publicly available reference cell profiles as surrogates (FS infloci, as suggested in the EDec method, see Material and Methods for probe selection information), we find a large reduction of error (47% error reduction on average with 20 patients and 26% with 100 patients). Then, we remove confounding probes, which corresponds to ∼1,000-2,000 probes significantly correlated with confounder variables (such as age, sex, etc.) with an adjusted false discovery rate (FDR) threshold of 15% (Benjamini and Hochberg 1995). We subsequently observe a substantial reduction of error (33% in average with 20 patients and 35% with 100 patients). Other FDR thresholds between 5% and 20% provide similar error measures (Supplementary Figure S6). After removal of correlated probes, there is no systematic advantage in filtering additional probes by feature selection (see Figure 3A and B, right panel). For each deconvolution software, best performances are always obtained, with both 20 and 100 patients, when removing probes correlated with confounders. However, the positive impact of removing confounding probes is only observed when inter-sample variation in mixture proportion is high (α0 = 1). When inter-sample variation is low (α0 = 10,000), the deconvolution is much more complicated, which could explain why we do not observe reproducible improvement of error detection after removing confounding probes in this case (Supplementary Figure S7).

**Figure 3:**
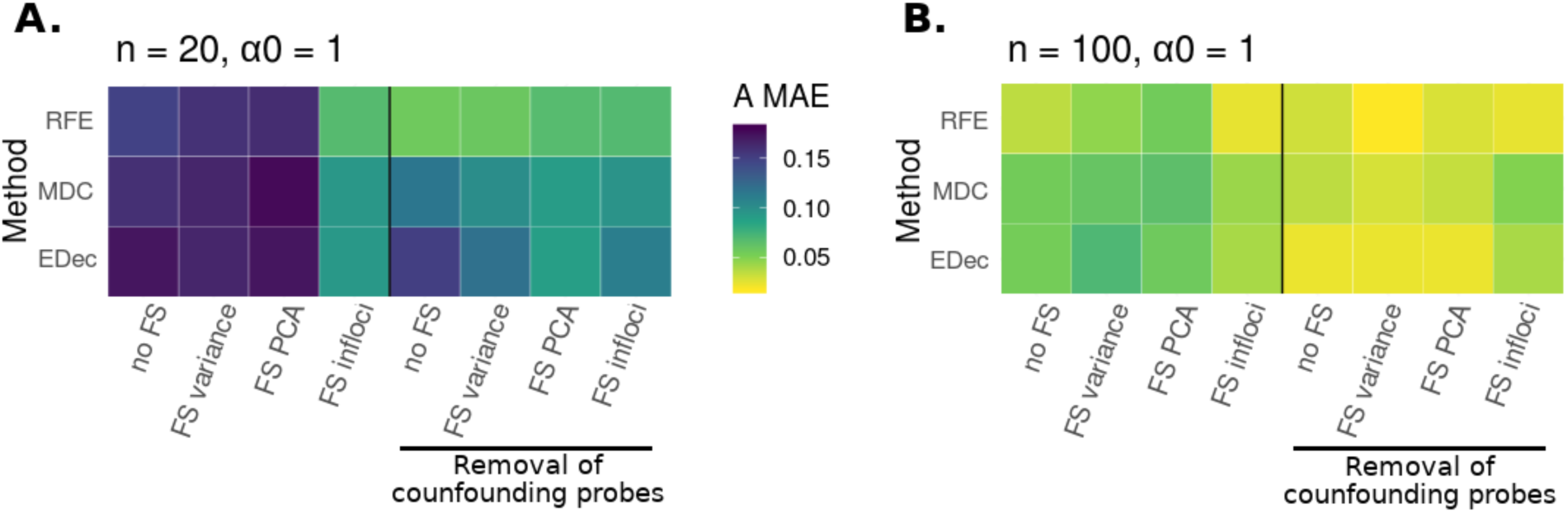
Impact of pre-treatments on method performance. Heatmap of method performances (‘A MAE’: Mean Absolute Error on estimated A, the matrix of cell proportions). RFE stands for RefFreeEWAS, MDC for MeDeCom and EDec for EDec stage 1. All algorithms are run on 10 D matrices: 10 different random noises *ε* were simulated on one matrix D computed from one simulated A matrix. In each heatmap, the left panel corresponds to algorithms run without accounting for confounders (no removal of confounding probes), the right panel corresponds to algorithms run accounting for confounders (removal of confounding probes by linear regression). In each case, different types of feature selection (FS) are tested: no FS = no feature selection, FS variance = selecting probes with high variance (var > 0.02), FS PCA = selecting probes highly correlated with the 4 first PCs (p_value < 0.1), FS infloci = selecting probes expected to biologically vary in methylation levels across constitutive cell types. (**A**) Simulations were performed with the following parameters K = 5, n = 20, α0 = 1, *ε* = 0.2 and G = 1. (**B**) Simulations were performed with the following parameters K = 5, n = 100, α0 = 1, *ε* = 0.2 and G = 1. The number of conserved probes is display supplementary table 1.

In summary, we highly recommend to systematically remove possible confounding probes to account for confounders. Filtering based on biologically informative loci, after removing of confounding probes, can also be an interesting approach, if the investigated biological system is properly defined. In the rest of the paper, removal of confounding probes is always considered before using MeDeCom, and RefFreeEwas whereas EDec infers informative probes from any available references (in EDec Stage 0, as also illustrated in its vignette).

### Choice of the number of cell types K

We tested several methods to choose the number of cell types K including the scree plot based on PCA of the D matrix, a cross validation score provided by MeDeCom, a deviance statistic estimated with bootstrap provided by RefFreeEWAS, and contrast those to the “best fit” and stability methods used by EDec.

First, we look at the scree plot by plotting the eigenvalues of the D matrix in descending order (Figure 4). For choosing K, we use Cattell’s rule, which states that components corresponding to eigenvalues to the left of the straight line should be retained (Cattell 1966). When the actual number of different cell types is equal to K, we expect that there are (K-1) eigenvalues would correspond to the mixture of cell types and that other eigenvalues would correspond to noise (or other unaccounted for confounders). Indeed, one PCA axis is needed to separate two types, two axes for three types, etc. However, when not accounting for confounders, Cattell’s rule overestimates the number of PCs and suggests to choose 3 and 5 PCs (i.e. K = 4 and K = 6) whereas the correct answers are K = 3 and K = 5 respectively (Figure 4 upper panel). When accounting for confounders by removing probes significantly correlated with confounders, Cattell’s rule provides the correct answer of 3 and 5 cell types (2 or 4 PCs) (Figure 4 lower panel). Finally, we observe that varying the FDR threshold used to select confounding probes has no impact of the choice of K using Cattell’s rule (for all thresholds tested, the estimation of K is correct, Supplementary Figure S8).

**Figure 4:**
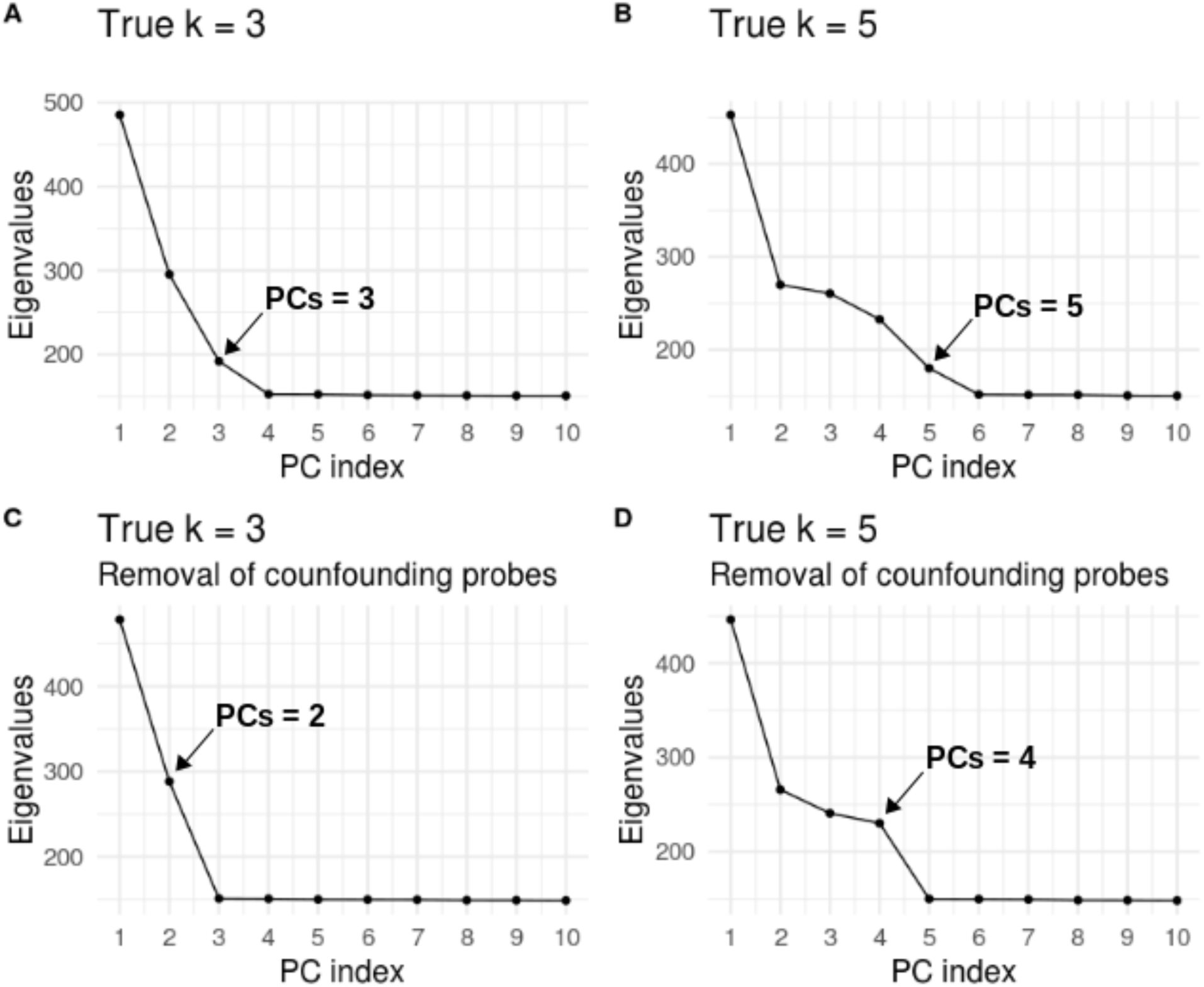
Determining K with PCA scree plot. To choose K, we recommend to use Cattell’s rule, calculating the estimated K as K = PCs + 1. The number of PCs chosen by the Cattel’s rule is shown with an arrow. The D matrix was simulated with the following parameters: n = 100, α0 = 1, *ε* = 0.2, G = 1 and K = 3 (**A** and **C**) and n = 100, α0 = 1, *ε* = 0.2, G = 1 and K = 5 (**B** and **D**). (**A, B**) Scree plot of PCA applied on D matrix before removal of confounding probes (23,381 probes). (**C, D**) Scree plot of PCA applied on D matrix after removal of confounding probes (22,551 probes in **C**, 22,532 probes in **D**).

The cross-validation score provided by MeDeCom gives similar choices of K as the scree plot. When not accounting for confounders, graphical inspection of the decay of cross-validation error, as a function of the number of cell types, suggests to keep K = 4 or K = 6 cell types. When accounting for confounders, it suggests to keep K = 3 or K = 5 cell types, which are the correct answers, yet the distinction is less obvious for K = 5 (Supplementary Figure S9 right panel).

The bootstrap estimation of the deviance provides different answers than the previous two approaches. When the exact value of the number of cell types is K = 3, the minimum value of deviance is reached at the correct value of K = 3, whether confounders were accounted for or not. When the exact value of the number of cell types is K = 5, the deviance statistic indicates to choose a K = 4 or 5 whether confounders were accounted for or not (Supplementary Figure S9 left panel).

Lastly, we investigate to what extent inference of cell type proportions is robust with respect to the choice of K. We find that the choice of K has a strong influence on inference of cell type proportions. Underestimation of K has a large impact on measured error, and overestimation of K, albeit to a lesser extent, also increases measured error (Figure 5). For instance, when the correct K is equal to 3, using K = 4 instead of K = 3 provides an error measure that is at least twice as large regardless of the deconvolution method that is used. Overestimation of K lead to large increase of error, even if our MAE-computing algorithm only retains the 3 cell types that minimize MAE error among the 4 inferred cell types.

**Figure 5:**
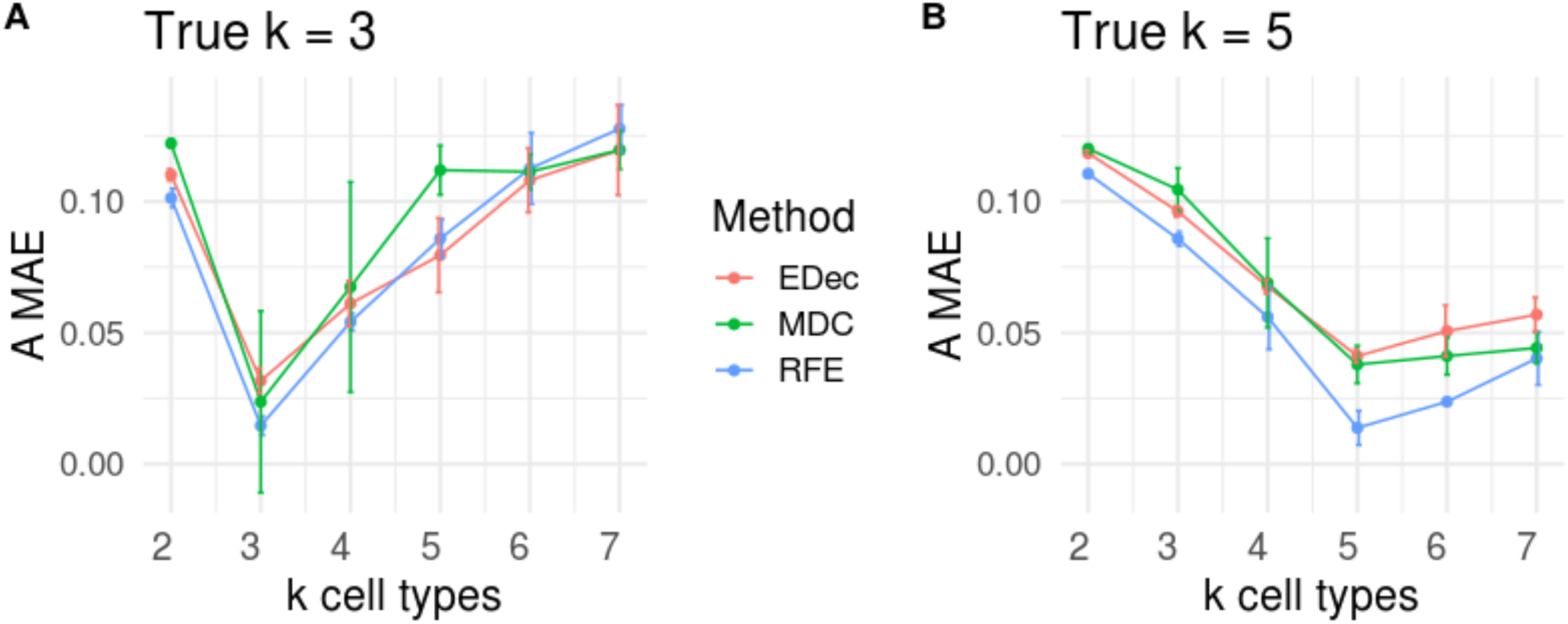
Impact of K selection on algorithm performance. ‘A MAE’ is shown for D matrices (mean value of 10 random noises applied on D) computed from 1 random A. Each color represents a different method. Error bars represent standard deviation on 10 random noise realizations. RFE stands for RefFreeEWAS, MDC for MeDeCom and EDec for EDec stage 1, each method was applied with various imposed K parameters (from 2 to 7). (**A**) The following parameters were used to simulate D : K =3, α0 = 1, *ε* = 0.2, G = 1 and n = 100. RFE and MDC methods were run after removal of confounding probes (between 22,517 and 22,624 remaining probes), EDec was run on informative loci, as recommended by the method’s authors (614 remaining probes). (**B**) The following parameters were used to simulate D : K =5, α0 = 1, *ε*= 0.2, G = 1 and n = 100. RFE and MDC methods were run after removal of confounding probes (between 22,551 and 22,602 remaining probes), EDec was run on informative loci, as recommended by the method’s authors (614 remaining probes).

These results suggest that the choice of K can be reliably guided by a scree plot based on PCA, after accounting for confounders.

Alternatively, the best performance at true K-s in Figure 5 suggests a different approach where K is selected after running deconvolution for various values of K and choosing the K with best performance. EDec adopts this general idea by selecting the K that explains the most variance (achieves the best fit of D) and largest K that shows stable estimates of both the A and T matrices. The stability is tested by taking at least 3 subsets (consisting of randomly selected 80% of the sample profiles) and measuring the similarity of estimates for A and for T across the 3 subsets.

### Biological interpretation of recovered methylation profiles T

We next compare the estimated matrix of average cell type-specific methylation profiles (matrix T) with the real methylation profiles of cell types used to simulate the datasets.

To assess the similarity of reconstructed methylation profiles, we run the three algorithms on a representative D matrix, with n = 100 patients. We then draw a heatmap representing the level of correlation between estimated methylation profiles and reference methylation profiles. When the inter-sample variation is high (α0 = 1), we observe that all the deconvolution methods tested succeed to properly estimate the cell type-specific methylation profiles and to robustly identify corresponding reference cell types (Figure 6). When the inter-sample variation is low (α0 = 10,000), all methods fail to identify reference cell types (Supplementary Figure S10), which is consistent with the results observed in Supplementary figure S7. Importantly, in a real setting, true methylation reference profiles are not available, and the identity of the cell-types present is unknown, which strongly complexifies the biological interpretation and the annotation of the obtained cell types.

**Figure 6:**
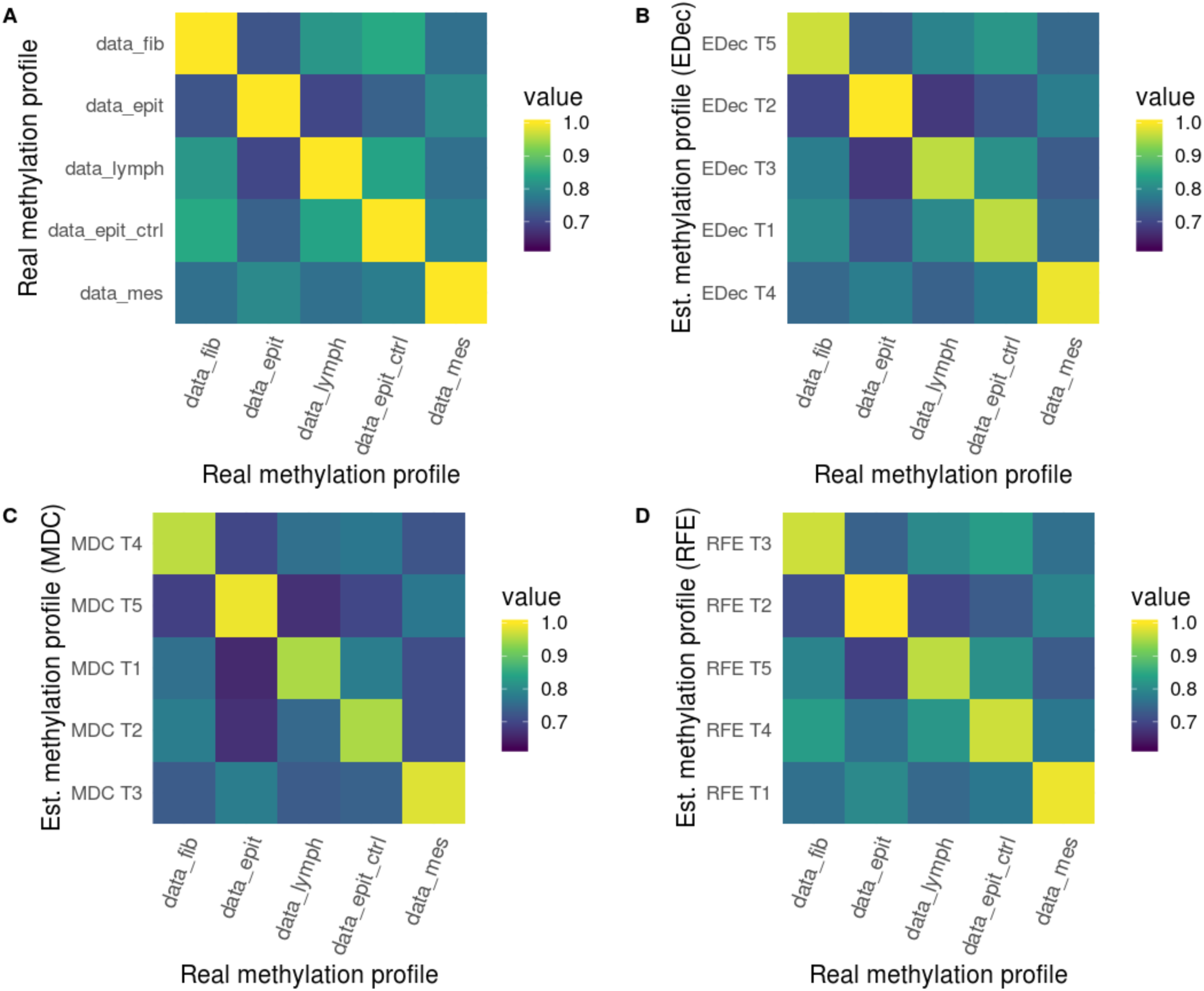
Correlation between estimated and real cell type-specific methylation profiles. Heatmap of the correlations between cell type-specific methylation profile used for the simulation and cell type-specific methylation profiles estimated (Est.) by different methods. In (**A**), the correlation between different cell types used for the simulation of the T matrix (data_fib = fibroblast, data_epith = cancerous epithelial, data_lymph = T lymphocytes, data_epit_ctrl = healthy epithelial and data_mes = cancerous mesenchymal). We applied EDec (**B**), MeDeCom (**C**) and RefFreeEwas (**D**) on a representative simulation of 100 patients (α0 = 1, *ε* = 0.2, G = 1, K = 5) after the removal of confounding probes by linear regression (22,483 remaining probes). We used the Pearson method to compute the correlation between the estimated cell type-specific methylation profiles and real cell type-specific methylation profiles used for the simulation.

Altogether, this indicates that reducing the inter-sample variability will impact both identification of cell type proportion and the identification of existing cell type-specific methylation profiles.

## Conclusion and discussion

### Recommendation

We have conducted extensive benchmarking of the core deconvolution step shared by three algorithms dedicated to reference-free cell-type heterogeneity quantification from DNA methylation datasets. We identify the following critical steps influencing method performance: i) accounting for confounders, ii) feature selection and iii) the choice of the number of estimated cell types. To account for confounders, we suggest to remove probes associated with the measured confounders. We suggest to use a large false discovery rate threshold, preferring to remove too many probes rather than keeping probes influenced by confounders, given the large (>20,000) initial number of probes. When probes associated with confounders are not removed, the number of cell types is overestimated, capturing the additional dimension of confounders (Figure 4). We recommend that care to be taken to choose the right number K of cell types, as the outcome of deconvolution may strongly depend on K. One such method is the scree plot and Cattell’s rule, which states that eigenvalues corresponding to true signal are to the left of the straight line (Figure 4). The number of cell types to consider is equal to the number of eigenvalues to keep plus one. Choice based on Cattell’s rule is robust with respect to the false discovery rate threshold used when discarding probes associated with confounders (Figure S7). Lastly, we suggest to use feature selection and to further reduce the number of probes by focusing on probes that are informative of cell types or the highest variance markers if no biological information is available. Further reducing the number of probes marginally affects estimation error (Figure 3), but can substantially reduce computational running time (Supplementary table 3). Once the markers and the number of cell types have been chosen, the core deconvolution step implemented by one of the three packages would provide comparable estimates of average methylation profiles (Figure 6) and of sample-specific cell-type proportions (Figure 7).

**Figure 7:**
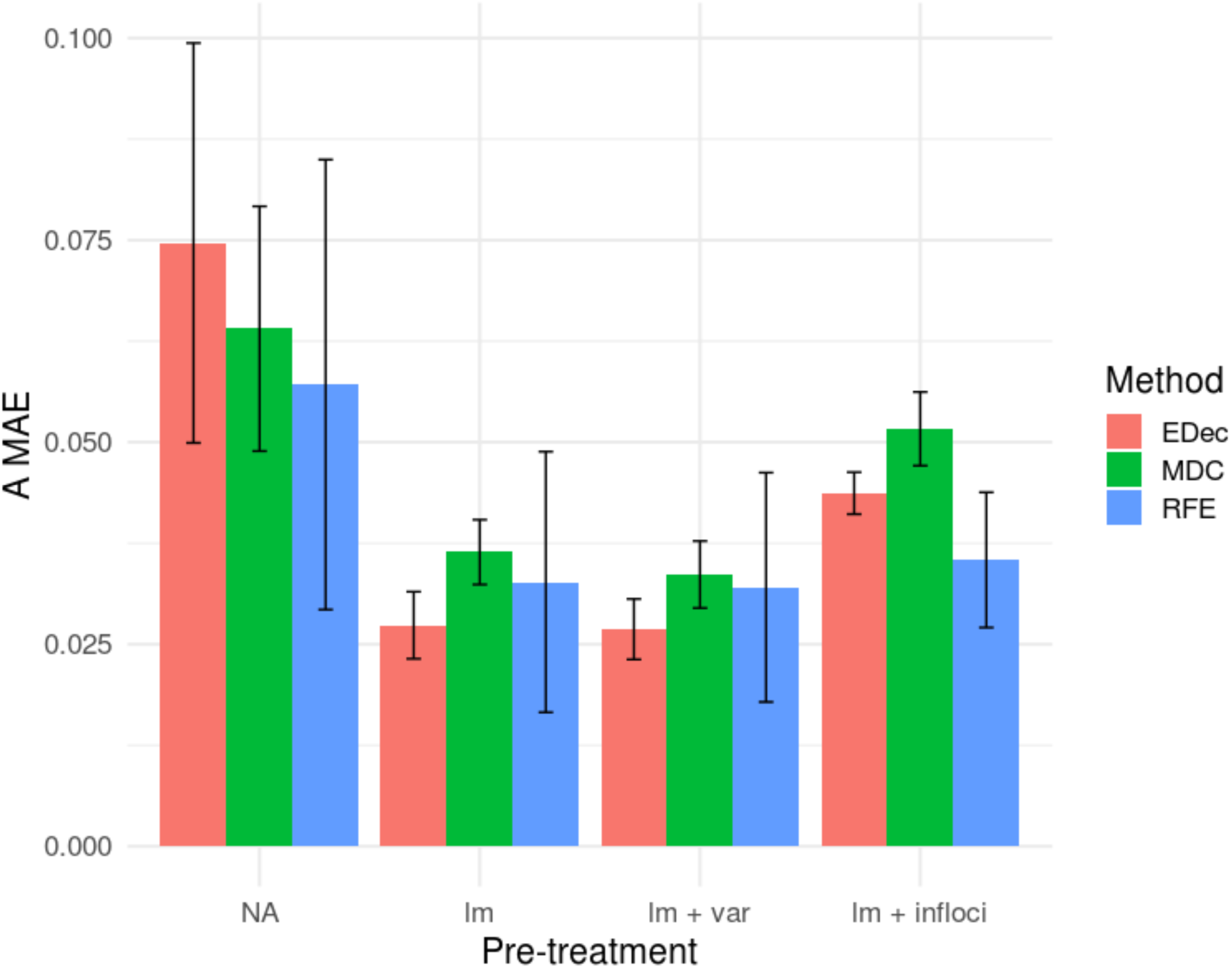
Comprehensive comparison of the pre-treatment pipeline. Histogram of ‘A MAE’ (‘A MAE’: Mean Absolute Error on estimated A, the matrix of cell proportions) for 10 D matrices (mean value of 10 random noises applied on D) computed from 10 random A. Each colour represents a different method. Error bars represent standard deviation on 10 random noises. RFE stands for RefFreeEWAS, MDC for MeDeCom and EDec for EDec stage 1. The following parameters were used to simulate D : K = 5, α0 = 1, *ε* = 0.2, G = 1. The methods were run without pre-treatment (NA), after removal of confounding probes by linear regression (lm), after removal of confounding probes and filtering for the most variable probes (lm + var), and after removal of confounding probes and filtering of probes expected to biologically vary in methylation levels across constitutive cell types (lm + infloci).

### Benchmark Pipeline

Based on lessons learned from the simulation experiments, we developed a benchmark pipeline to estimate cell-type proportions that addresses the presence of confounders and other key factors affecting performance of deconvolution algorithms (Figure 8). We anticipate that this benchmark pipeline will help catalyze wide adoption of deconvolution methods and accelerate improvement of deconvolution pipelines by (1) helping validate other deconvolution pipelines by demonstrating concordant results; (2) serving as a benchmark for demonstrating improved performance of other pipelines; (3) providing a starting point (“toolkit”) for development of new pipelines.

**Figure 8:**
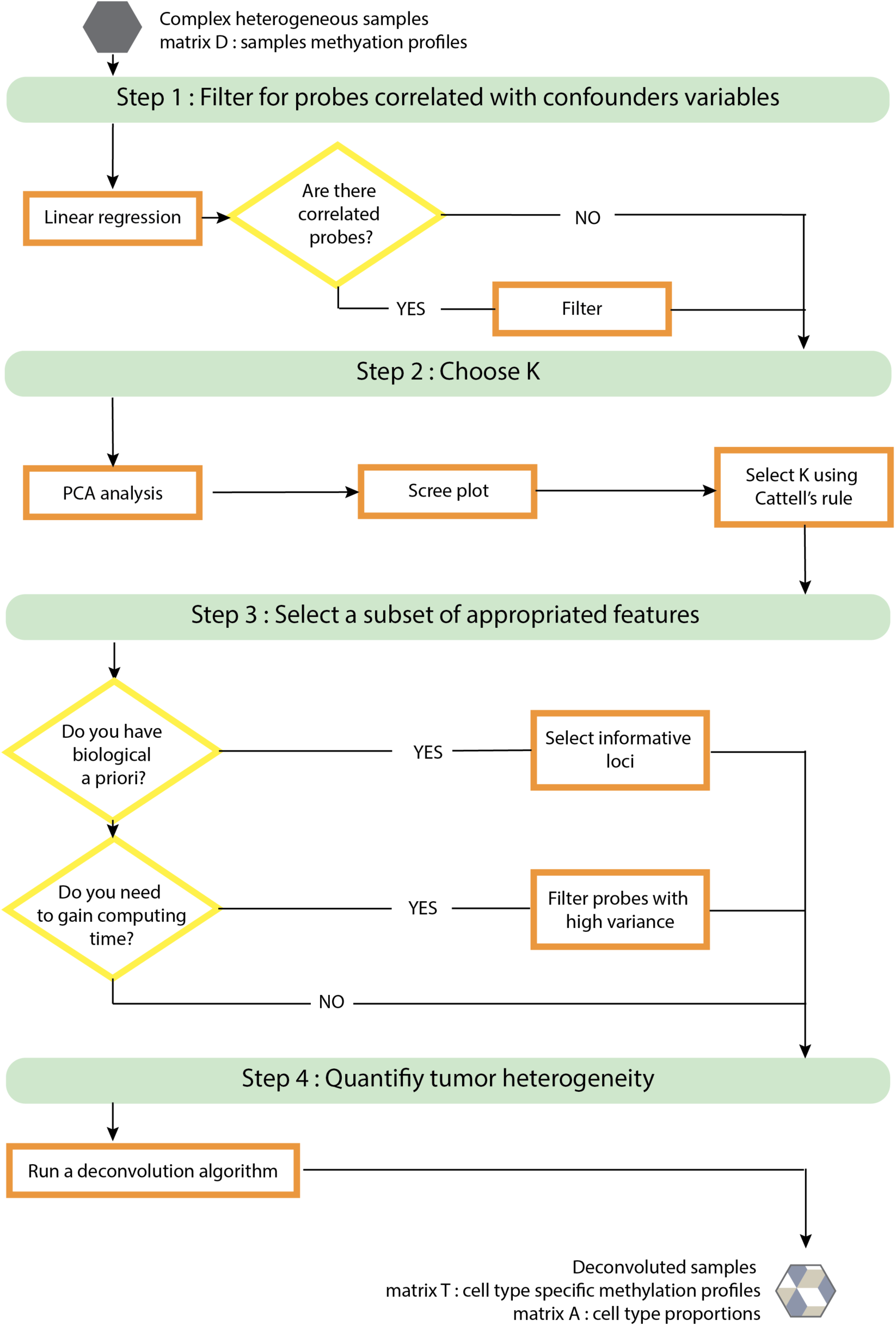
Recommendations and benchmarking pipeline.

We note that the benchmark pipeline is not experimentally validated nor it is systematically compared as a whole against more complex pipelines that include expression data (e.g., all stages of EDec pipeline). In our experience, no deconvolution pipeline can be expected to provide accurate solutions when applied “out of the box” to a new tumor type. Tuning and validation are required in the context of each tumor type, using resources and information that may be tumor-type specific. In that sense, deconvolution may be thought of as a computational modeling approach that goes hand-in-hand with experimentation.

### Initialization

The software MeDeCom, EDec and RefFreeEwas have different methods to initialize the deconvolution algorithm. MeDeCom performs multiple random initialization of the matrix of proportion A. EDec performs initialization with random draws of proportions of cell types in each sample. RefFreeEwas initializes the matrix of cell types using reduction dimension or clustering techniques depending on a user-defined option. For RefFreeEwas, we find that clustering techniques provide the best option in the favorable scenario when cell-type proportions strongly differ between individuals (Figure 2). However, in the less favorable scenario where cell-type proportions are more similar between individuals, initialization based on singular value decomposition should be preferred (Figure S4). Differences of performance according to initialization are substantial and depending on initialization, error measures can vary by a factor of two. The fact that deconvolution methods depend on initialization indicates that there is room for improvement either by finding an optimal initialization strategy or by using an ensemble method that combines or averages several deconvolution solutions (Alhamdoosh et al. 2017; Jakobsson and Rosenberg 2007).

### Evaluation of performance

To estimate performances of methods, we used the MAE metric. A similar alternative metric, the root mean square error (RMSE) gave equivalent performance evaluation (Supplementary Figure S11). Internal steps of the MeDeCom algorithm use a leaving-columns-out Cross-Validation Error (CVE) to choose regularization parameter λ and number of cell types K. In our study, we consider a grid of six values for the regularization parameter λ (0, 0.00001, 0.0001, 0.001, 0.01, 0.1). Interestingly, the optimum lambda selected by the CVE approach does not always perform better than λ = 0, when evaluated by the MAE metric (Supplementary Figure S12). Although it is difficult to assess the biological relevance of each possible error metric, we would like to emphasize that the choice of evaluation metrics is an important parameter when conducting a benchmarking study. In addition, evaluation based on simulations remain limited compared to evaluation based on real tumor datasets, because the *in silico* simulation does not model all the biological properties of the system, such as changes in methylation due to cell-cell interaction. However, we are still lacking real tumor dataset with accurate quantification of tumor heterogeneity. Deconvolution algorithms evaluation will then be significantly improved with the generation of dedicated *in vivo* benchmarking dataset

### Data challenge, collaborative and open science

Our work strongly benefits from a data challenge format where different pipelines were proposed and evaluated. We gathered methylation deconvolution experts for a week of brainstorming on this benchmarking issue. We used the resulting ideas and computational methods to construct several pipelines which we evaluated in the paper. Thus, all challenge participants are referred as consortium authors of the paper. As a key deliverable of the project, to facilitate wide application of reference-free deconvolution and also pipeline development by the community, we develop a benchmark pipeline and release it as an R package (Figure 8 and R package *medepir*).

## Material and methods

### Matrix factorisation

We assume D is a (M×N) methylation matrix composed of methylation value for N samples, at M CpG methylation sites. Each sample is constituted of K cell types. We assume the following model: D = TA, with T an unknown (M×K) matrix of K cell type-specific methylation reference profiles (composed of M sites), and A an unknown (K×N) proportion matrix composed of K cell type proportions for each sample. In the methods tested here, A and T are found using matrix factorization, which consists of minimizing the error term ||*D* – *TA*||_2_, with constraints on methylation values, 0 ≤ *A* ≤ 1 and 0 ≤ *T* ≤ 1, and on proportions 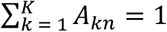(MeDeCom and EDec) or 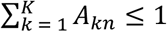(RefFreeEWAS), where *A* _*kn*_is the proportion of the n*th* sample for the k*th* cell type. MeDeCom uses an additional regularization function, which depends on a regularization term, which is weighted by a hyperparameter λ, that favors methylation values close to 0 or 1 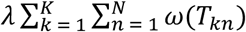 with *ω*(*x*)=*x(1-x*)

### Simulations

We used the The Cancer Genome Atlas (TCGA) lung cancer dataset (LUAD and LUSC) as an example biological dataset. We simulated synthetic DNA methylation mixtures. We selected M = 23,381 probes using the following criteria: (i) intersect between the Illumina 27k and Illumina 450k DNA methylation array, and (ii) non-null probes in 100% of LUAD and LUSC methylation datasets. Simulated D matrix were obtained using D = TA model.

The A matrix is simulated by a Dirichlet distribution with parameters defined for 10% of fibroblast, 60% of cancerous epithelial, 5% of T lymphocytes, 15% of control epithelial and 10% of cancerous mesenchyme. The mixture proportions are sampled from a Dirichlet distribution with parameters generating sets of proportions more or less variable across the sample population. The parameter *α*0, which defines the variability across the sample population, is set to 1 by default. For simulation with only three cell types, we used 20% of fibroblast, 70% of cancerous epithelial and 10% of T lymphocytes.

To simulate the T matrix of cell type-specific methylation profile, we used different study from GEO. For the cancerous epithelial line, we used the cell line GSM1560930 and for the cancerous mesenchymal cell line, we used the cell line GSM1560925, from the same study. For the fibroblast cell line, we used the cell line GSM1354676 after conversion from m-value to b-value. For the T lymphocyte cell line, we used the GSM1641099 and for the control epithelial the GSM2743808. These cell lines correspond to the cell line background G1. To simulate the G2, we used different cell line from the same studies. The cell line of cancerous epithelial is GSM1560911, these of the cancer mesenchymal is GSM1560931, the fibroblast cell line is GSM1354675, the T cell line is GSM1641101 and the control epithelial cell line is GSM2743807.

Once the initial T matrix is defined, we applied a series of confounding effects using parameter extracted from the clinical data associated with LUAD and LUSC cohorts.

First, we generate an individual-specific T matrix, accounting for two major biological confounders for methylation, which are sex and age.

To identify effects of sex, we performed linear regression of methylation by sex in the TCGA dataset and detected 1397 probes correlated with sex (p-value < 0.01). Given that the majority of cell lines used to construct the initial T matrix were derived from male individuals, we used the corresponding linear regression coefficients to shift accordingly methylation value of female-associated T matrices. We used the same sex coefficient for each cell type represented in the T matrix.

To identify effects of age, we performed linear regression of methylation by age in the TCGA dataset and identified 113 probes correlated with age (p-value < 0.05). We used this linear model to generate an individual-specific methylation profile, according to its age. Then, we arbitrarily decided to assign these 113 methylation values to the normal epithelial cell type. For each probe associated with age, we then applied a normalization coefficient (ratio of each cell type to the epithelial cell type computed in the initial T matrix) to modify methylation values of the remaining cell types. Our simulation scheme implicitly assumes that age has the same effect whatever the cell type.

Second, we decided to account for technical confounders. We calculated 22 median plate-effects (TCGA experimental batch effect) using 1000 random probes. For each probe, we modeled plate effects using multiplicative coefficients that measure the ratio of mean methylation values of a plate on mean methylation of the (arbitrarily) 1st plate. Each coefficient is estimated by the median of the 1000 ratios of methylation values. These multiplicative coefficients are then used on all probes to model batch effects on the matrix D of individual convoluted methylation profile.

Finally, we added gaussian noise on the matrix of convoluted methylation profiles D. By default, we used the gaussian parameters mean = 0 and sd = 0.2. In case noise generated methylation values larger than 1 or smaller than 0, noise was not added to the methylation value.

The simulation function is accessible using our R package medepir.

### Software usage

We follow publication guidelines and default parameters for each method (see Material and Methods for details in software usage). RefFreeEwas was used with the function “RefFreeCellMix” with 9 iterations and remaining parameters set to default, unless specified otherwise. MeDeCom was used with the function runMeDeCom with the following parameters: NINIT = 10, NFOLDS = 10, ITERMAX = 300, lambdas=c(0, 0.00001, 0.0001, 0.001, 0.01, 0.1), if unless specified otherwise. EDec was used with the function run_edec_stage_1 with default parameters, unless specified otherwise. When testing the initialization of matrix T in RefFreeEWAS algorithm, we used the following functions: RefFreeEWAS::RefFreeCellMixInitializeBySVD(D, type = 1), RefFreeEWAS:: RefFreeCellMixInitialize(method = “ward”, dist.method = “euclidean”), RefFreeEWAS:: RefFreeCellMixInitialize(method = “ward”, dist.method = “manhattan”). Each RefFreeEWAS initialization method estimates an averaged methylation profile for K components from the initial D matrix (either by hierarchical clustering on D, or by singular value decomposition of D).

### Confounding factor detection

Accounting for confounding probes was performed using linear regression for each confounding factor (based on clinical metadata extracted from TCGA LUAD and LUSC cohorts). We control for false discovery rate using Benjamini-Hochberg correction. For each confounder variable, we removed probes that are significantly associated with the confounder variable using a false discovery rate threshold of 0.15.

### Feature selection

To enhance the precision of the method, we can select the more informative probes. For this, we tested three methods: (i) selecting probes with high variance (var > 0.02), (ii) selecting probes highly correlated with the first four PCs (p_value < 0.1) (Luu, Bazin, and Blum 2017; Prive, Aschard, and Blum, n.d.), and (iii) selecting probes expected to biologically vary in methylation levels across constitutive cell types, as described in EDec method(Prive, Aschard, and Blum, n.d.; Luu, Bazin, and Blum 2017), the corresponding probes are depicted in supplementary table S4.

### Performance evaluation

We evaluate algorithm performances by (i) computing the mean absolute error (MAE) on estimated A, defined as 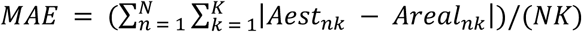, or (ii) computing the root-mean-square error (RMSE) on estimated A, defined as 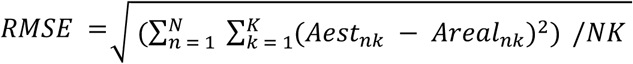, with n the total number of observations.

## Declarations

### Ethics approval and consent to participate

Not applicable.

### Consent for publication

Not applicable.

### Availability of data and material

The *medepir* R package (DNA MEthylation DEconvolution PIpeline in R) can be downloaded at https://github.com/bcm-uga/medepir. Documentation and usage examples are also available on the same page. All the datasets associated with this publication (lung cancer patient metadata) can be found in the TCGA webpage.

### Author’s contributions

MR conceived and designed the project. MR, CD, FP, AW and RB developed the method. CD and FP implemented the R package. MR, CD, FP, and MB analyzed the results. MR and MB wrote the manuscript with the help of CD, FP, DJ, EL, PL, EAH PL, AM and MS. All authors read and approved the final manuscript.

## Acknowledgements

We thank the members of the BCM team for inspiring discussions during regular joint group meetings. We are grateful to the Codalab data challenge open source platform. We thank all members of the HADACA consortium for helpful discussion and contributions during the methylation deconvolution data challenge (December 2018, Aussois, France).

## Supplementary material

**Supplementary Figures**: **Figure S1**: Overview of *in silico* simulations. **Figure S2**: Variation of the estimated A MAE depending of the number of patients. **Figure S3**: Distribution of cell types proportions according to the parameter α0. **Figure S4**: Impact of the initialisation method of RefFreeEwas for a stable Dirichlet simulation. **Figure S5**: Effect of A matrix initialization on algorithms performances. **Figure S6**: Impact of the FDR threshold for the removing of confounding factors. **Figure S7**: Impact of pre-treatments for a stable dirichlet. **Figure S8**: Determining K is robust to variations in accounting for confounders. **Figure S9**: Determining K by RefFreeEWAS and MeDeCom. **Figure S10**: Correlation between estimated and real cell type-specific methylation profiles for a Dirichlet of α0 = 10,000. **Figure S11**: Variation of the error metric: Mean Absolute Error and Root-Mean-Square Error. **Figure S12**: Impact of lambda parameter for MeDeCom.

**Supplementary Tables: Supplementary Table 1**: Number of remaining probes in figure 2. **Supplementary Table 2**: Number of remaining probes in figure S3. **Supplementary Table 3**: Mean execution time in figure 7 (minutes).

## References

Alhamdoosh, Monther, Milica Ng, Nicholas J. Wilson, Julie M. Sheridan, Huy Huynh, Michael J. Wilson, and Matthew E. Ritchie. 2017. “Combining Multiple Tools Outperforms Individual Methods in Gene Set Enrichment Analyses.” Bioinformatics 33 (3): 414–24.

Alizadeh, Ash A., Victoria Aranda, Alberto Bardelli, Cedric Blanpain, Christoph Bock, Christine Borowski, Carlos Caldas, et al. 2015. “Toward Understanding and Exploiting Tumor Heterogeneity.” Nature Medicine 21 (8): 846–53.

Benjamini, Yoav, and Yosef Hochberg. 1995. “Controlling the False Discovery Rate: A Practical and Powerful Approach to Multiple Testing.” Journal of the Royal Statistical Society: Series B (Methodological). https://doi.org/10.1111/j.2517-6161.1995.tb02031.x.

Cattell, R. B. 1966. “The Scree Test For The Number Of Factors.” Multivariate Behavioral Research 1 (2): 245–76.

Houseman, E. Andres, Molly L. Kile, David C. Christiani, Tan A. Ince, Karl T. Kelsey, and Carmen J. Marsit. 2016. “Reference-Free Deconvolution of DNA Methylation Data and Mediation by Cell Composition Effects.” BMC Bioinformatics 17 (June): 259.

Jakobsson, Mattias, and Noah A. Rosenberg. 2007. “CLUMPP: A Cluster Matching and Permutation Program for Dealing with Label Switching and Multimodality in Analysis of Population Structure.” Bioinformatics 23 (14): 1801–6.

Jones, Peter A. 2012. “Functions of DNA Methylation: Islands, Start Sites, Gene Bodies and beyond.” Nature Reviews. Genetics 13 (7): 484–92.

Kaushal, Akhilesh, Hongmei Zhang, Wilfried J. J. Karmaus, Meredith Ray, Mylin A. Torres, Alicia K. Smith, and Shu-Li Wang. 2017. “Comparison of Different Cell Type Correction Methods for Genome-Scale Epigenetics Studies.” BMC Bioinformatics 18 (1): 216.

Lutsik, Pavlo, Martin Slawski, Gilles Gasparoni, Nikita Vedeneev, Matthias Hein, and Jörn Walter. 2017. “MeDeCom: Discovery and Quantification of Latent Components of Heterogeneous Methylomes.” Genome Biology 18 (1): 55.

Luu, Keurcien, Eric Bazin, and Michael G. B. Blum. 2017. “Pcadapt: An R Package to Perform Genome Scans for Selection Based on Principal Component Analysis.” Molecular Ecology Resources 17 (1): 67–77.

McGregor, Kevin, Sasha Bernatsky, Ines Colmegna, Marie Hudson, Tomi Pastinen, Aurélie Labbe, and Celia M. T. Greenwood. 2016. “An Evaluation of Methods Correcting for Cell-Type Heterogeneity in DNA Methylation Studies.” Genome Biology 17 (May): 84.

Onuchic, Vitor, Ryan J. Hartmaier, David N. Boone, Michael L. Samuels, Ronak Y. Patel, Wendy M. White, Vesna D. Garovic, et al. 2016. “Epigenomic Deconvolution of Breast Tumors Reveals Metabolic Coupling between Constituent Cell Types.” Cell Reports 17 (8): 2075–86.

Prive, Florian, Hugues Aschard, and Michael G. B. Blum. n.d. “Efficient Management and Analysis of Large-Scale Genome-Wide Data with Two R Packages: Bigstatsr and Bigsnpr.” https://doi.org/10.1101/190926.

Roadmap Epigenomics Consortium, Anshul Kundaje, Wouter Meuleman, Jason Ernst, Misha Bilenky, Angela Yen, Alireza Heravi-Moussavi, et al. 2015. “Integrative Analysis of 111 Reference Human Epigenomes.” Nature 518 (7539): 317–30.

Teschendorff, Andrew E., Charles E. Breeze, Shijie C. Zheng, and Stephan Beck. 2017. “A Comparison of Reference-Based Algorithms for Correcting Cell-Type Heterogeneity in Epigenome-Wide Association Studies.” BMC Bioinformatics 18 (1): 105.

Titus, Alexander J., Rachel M. Gallimore, Lucas A. Salas, and Brock C. Christensen. 2017. “Cell-Type Deconvolution from DNA Methylation: A Review of Recent Applications.” Human Molecular Genetics. https://doi.org/10.1093/hmg/ddx275.

Zheng, Shijie C., Charles E. Breeze, Stephan Beck, and Andrew E. Teschendorff. 2018. “Identification of Differentially Methylated Cell Types in Epigenome-Wide Association Studies.” Nature Methods 15 (12): 1059–66.

