## supplementary_figures for "Guidelines for cell-type heterogeneity quantification based on a comparative analysis of reference-free DNA methylation deconvolution software"

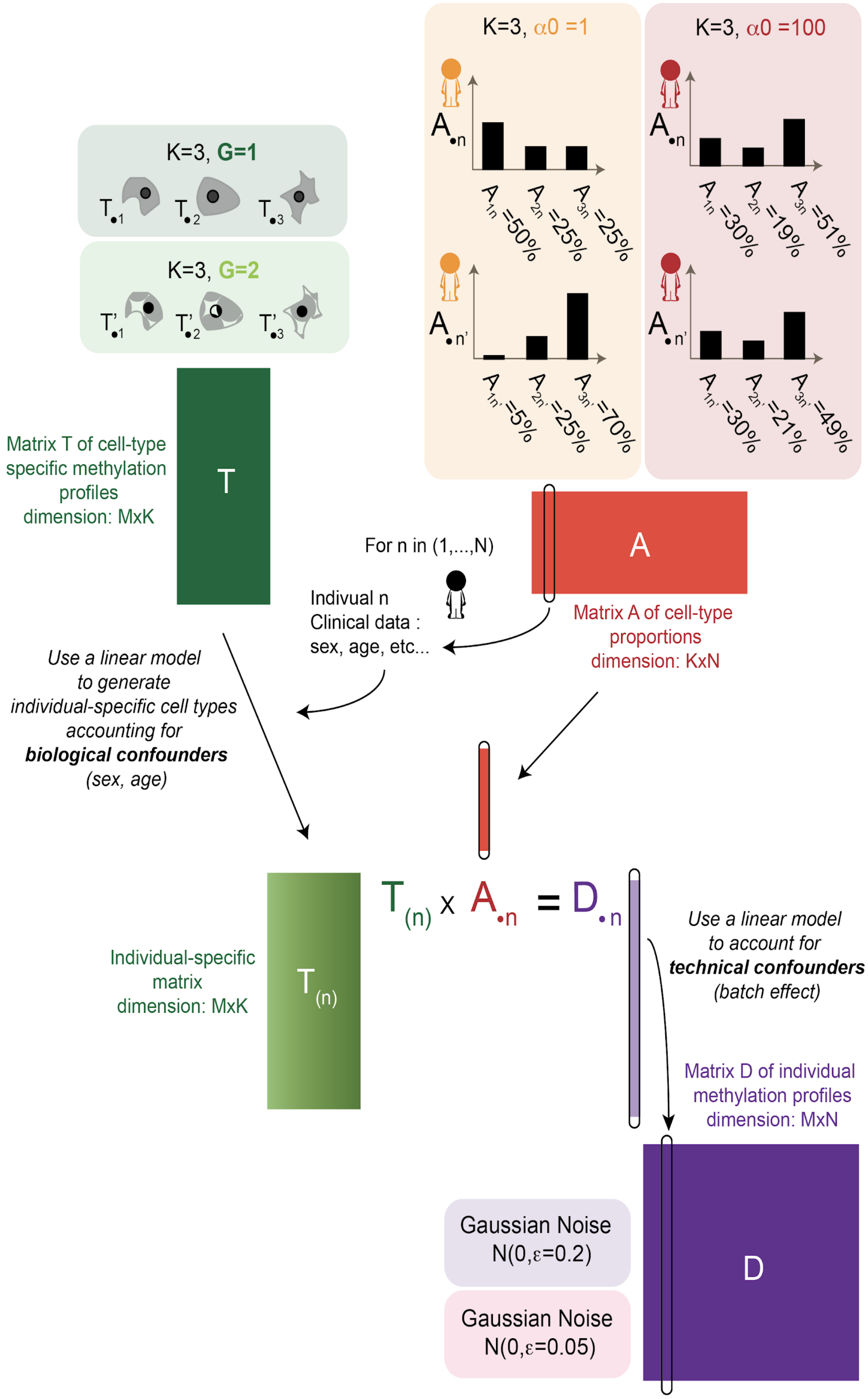


Figure S1: Overview of *in silico* simulations

Simulation of D matrix accounting for confounders effect. In our model, we simulate a individual-specific T(n) matrix of cell type-specific methylation profiles that account for sex and age of the individual n (with n ranging from 1 to N). Parameters N and α0 directly affect simulations of matrix A. Parameter G affects the construction of matrix T. Parameter $\varepsilon$ affects the final methylation values of matrix D.

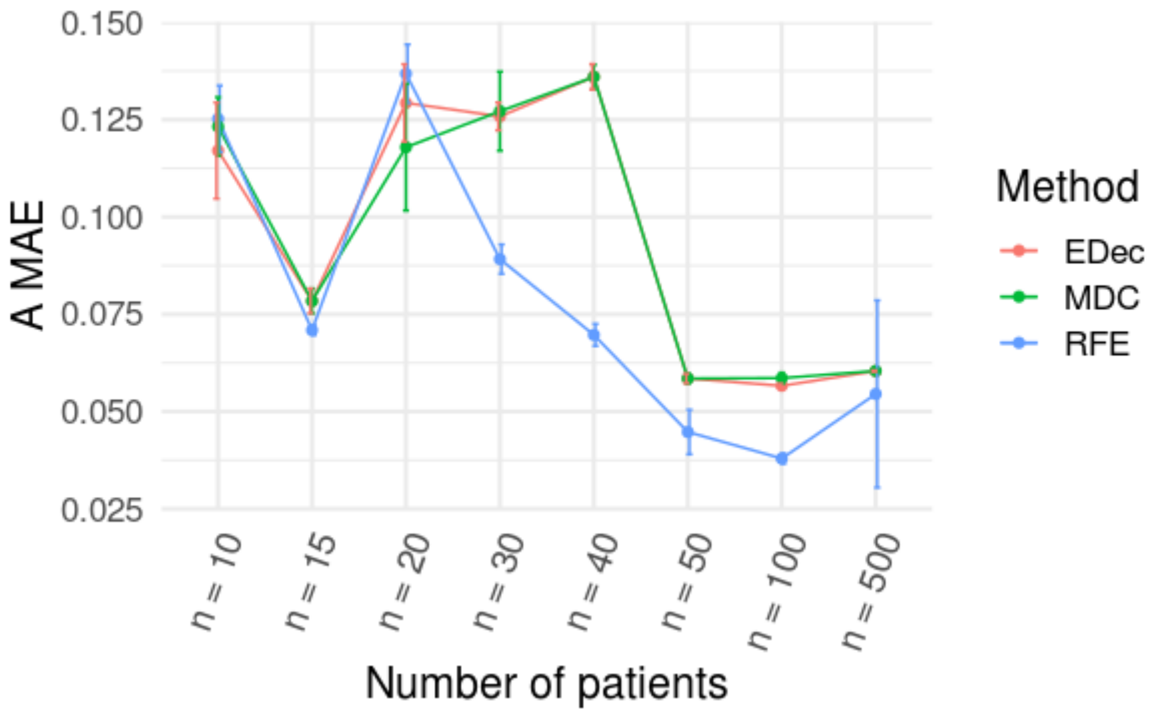


Figure S2: Variation of the estimated A MAE depending of the number of patients

In the figure 1, EDec, MeDeCom and RefFreeEWAS were run on 10 different random noise realisations for each parameter and the mean of these 10 error was represented. Here, the detail of A MAE values with the standard deviation is shown for the simulations of variation of the number of patients ($\alpha 0 = 1$, $\varepsilon = 0.2$, $G = 1$).

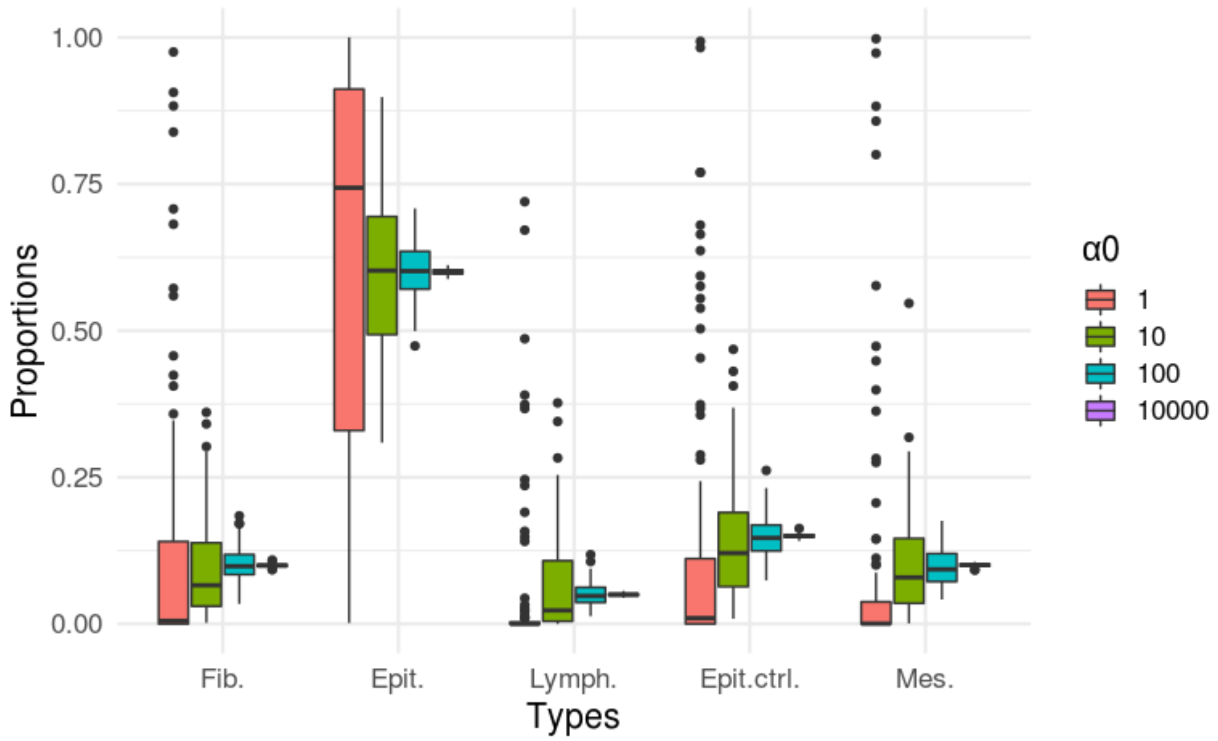


Figure S3: Distribution of cell types proportions according to the parameter α0

Boxplot of the proportions of different cell types simulated for 100 patients by a Dirichlet distribution of theoretical proportions: 10% fibroblast, 60% cancerous epithelial cells, 5% T lymphocytes, 15% normal epithelial cells and 10% cancerous mesenchymal cells. The α0 parameter influence the variability of proportions between the samples.

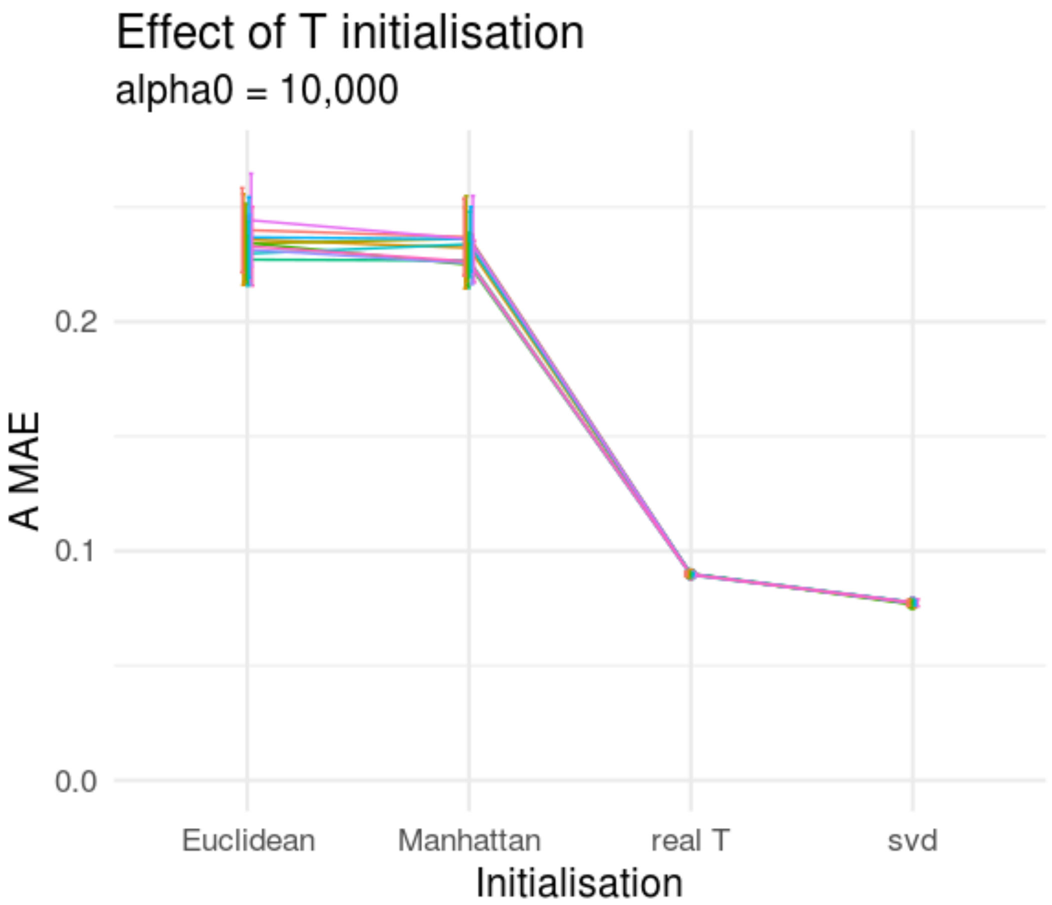


Figure S4: Impact of the initialisation method of RefFreeEwas for a stable Dirichlet simulation

`A MAE` is shown for 10 D matrices (mean value of 10 random noises applied on D) computed from 10 random A. Each colour represents a different simulated A. Error bars represent standard deviation on 10 random noises. The following parameters were used to simulate D: K =5, α0 = 10,000, $\varepsilon$ = 0.2, G = 1 and n = 100). Euclidean corresponds to RefFreeEWAS::RefFreeCellMixInitialize function applied with the default parameter dist.method = “euclidean”. Manhattan corresponds to RefFreeEWAS::RefFreeCellMixInitialize function applied with the parameter dist.method = “manhattan”. Real T corresponds to RefFreeEWAS::RefFreeCellMix used with the parameter mu0 = real_T, with real_T the matrix composed of the 5 cell types used to simulate D. svd corresponds to RefFreeEWAS::RefFreeCellMixInitializeBySVD function with default parameters.

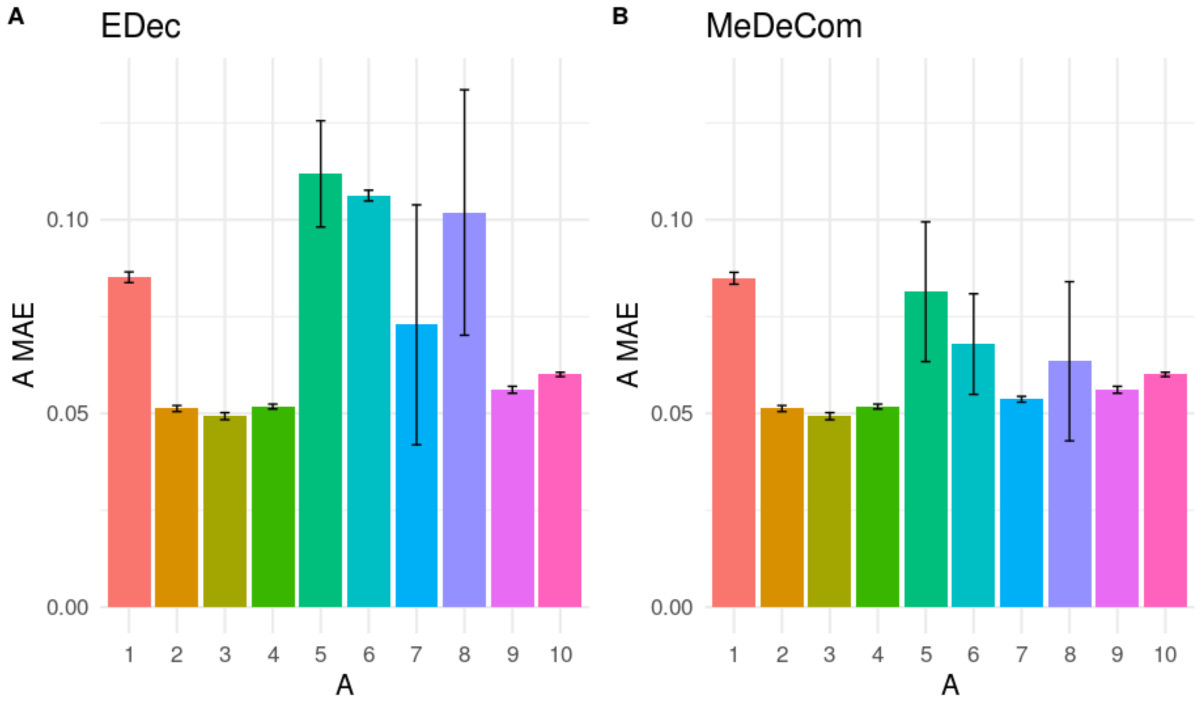


Figure S5 : Effect of A matrix initialization on algorithms performances

A MAE is shown for 10 D matrices generated with 10 different random A matrices with the same parameter α0 = 1. Error bars represent standard deviation on 10 random noises. The following parameters were used to simulate D: $\varepsilon$ = 0.2, G = 1 and n = 100. EDec (A) and MeDeCom (B) was run on the 100 matrices. To compare EDec and MeDeCom initialization approaches, MeDeCom was run with the regularization parameter $\lambda = 0$

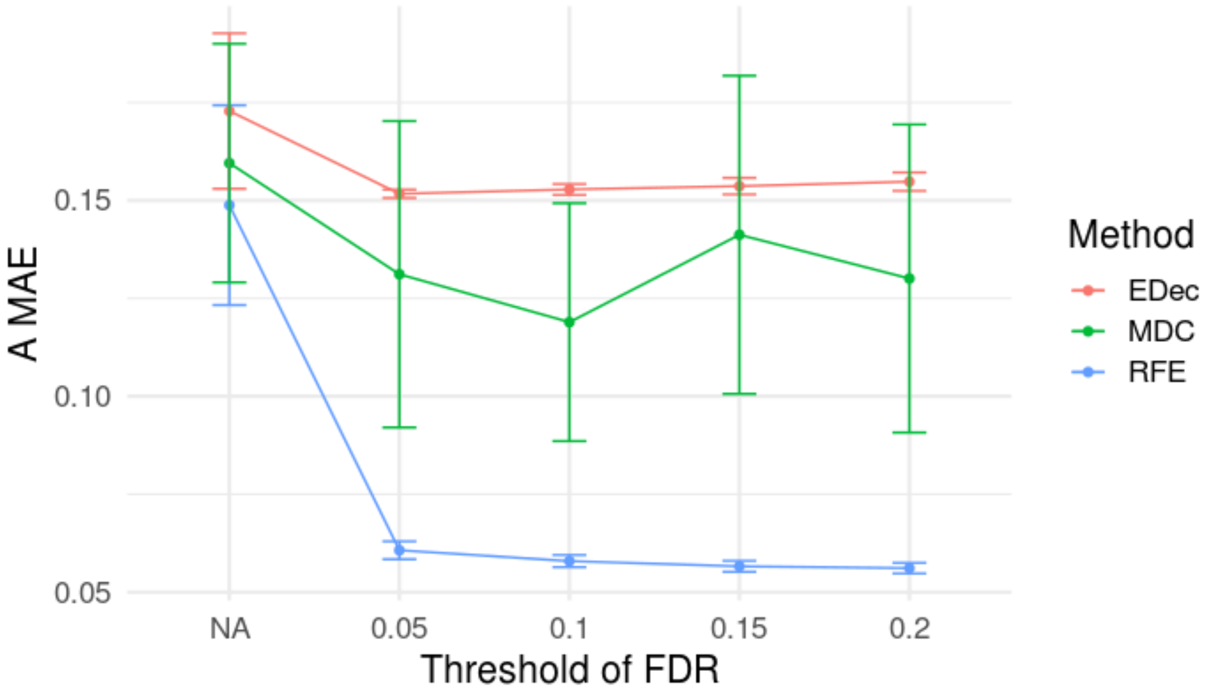


Figure S6: Impact of the FDR threshold for the removing of confounding factors

Confounding probes are removed by linear model, the FDR (False Discovery Rate) threshold impact the number of probes removed. The `A MAE` (Mean Absolute Error) was computed for different threshold and for 10 random noises applied on a D matrix. NA correspond to the D matrix before pre-treatments. Error bars represent standard deviation on 10 random noises. The following parameters were used to simulate D: K = 5, $\alpha 0$ = 1, $\varepsilon$ = 0.2, G = 1 and n = 20). For a threshold of 0.05, we keep on average 22,540 probes, for threshold = 0.1 we keep on average 21,998 probes, for threshold = 0.15 we keep on average 21,489 and for threshold = 0.2 we keep on average 20,950.

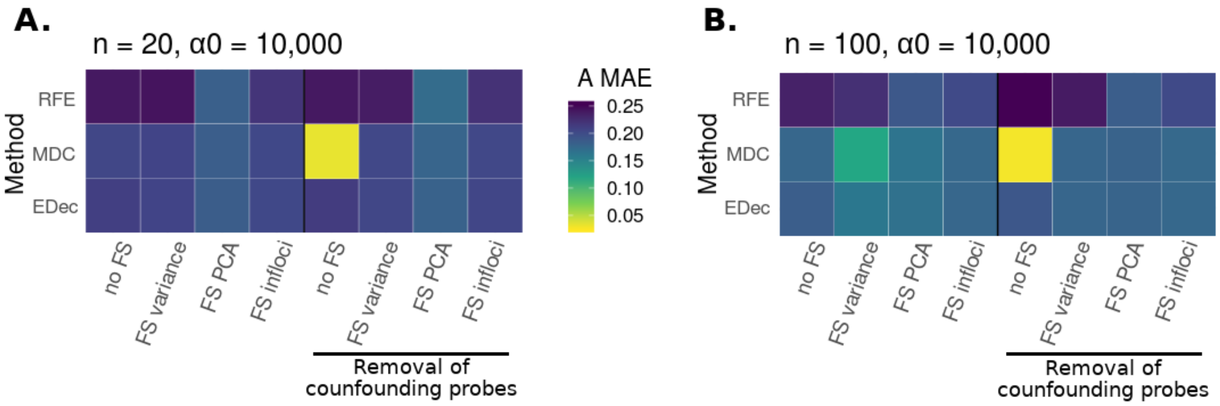


Figure S7: Impact of pre-treatments for a stable dirichlet

Heatmap of method performances (`A MAE`: Mean Absolute Error on estimated A, the matrix of cell proportions). RFE stands for RefFreeEWAS, MDC for MeDeCom and EDec for EDec stage 1. All algorithms were run on 10 D matrices: 10 different random noises $\varepsilon$ were simulated on one matrix D computed from one simulated A matrix. In each heatmap, the left panel corresponds to algorithms run without accounting for confounders (no removal of confounding probes), the right panel corresponds to algorithms run accounting for confounders (removal of confounding probes by linear regression). In each case, different types of feature selection (FS) are tested: no FS = no feature selection, FS variance = selecting probes with high variance (var > 0.02), FS PCA = selecting probes highly correlated with the 4 first PCs (p_value < 0.1), FS infloci = selecting probes expected to vary in methylation levels across constitutive cell types. (**A**) Simulations were performed with the following parameters K = 5, n = 20, α0 = 10,000, $\varepsilon$ = 0.2 and G = 1. (**B**) Simulations were performed with the following parameters K = 5, n = 100, α0 = 10,000, $\varepsilon$ = 0.2 and G = 1. The number of conserved probes is display supplementary table 2.

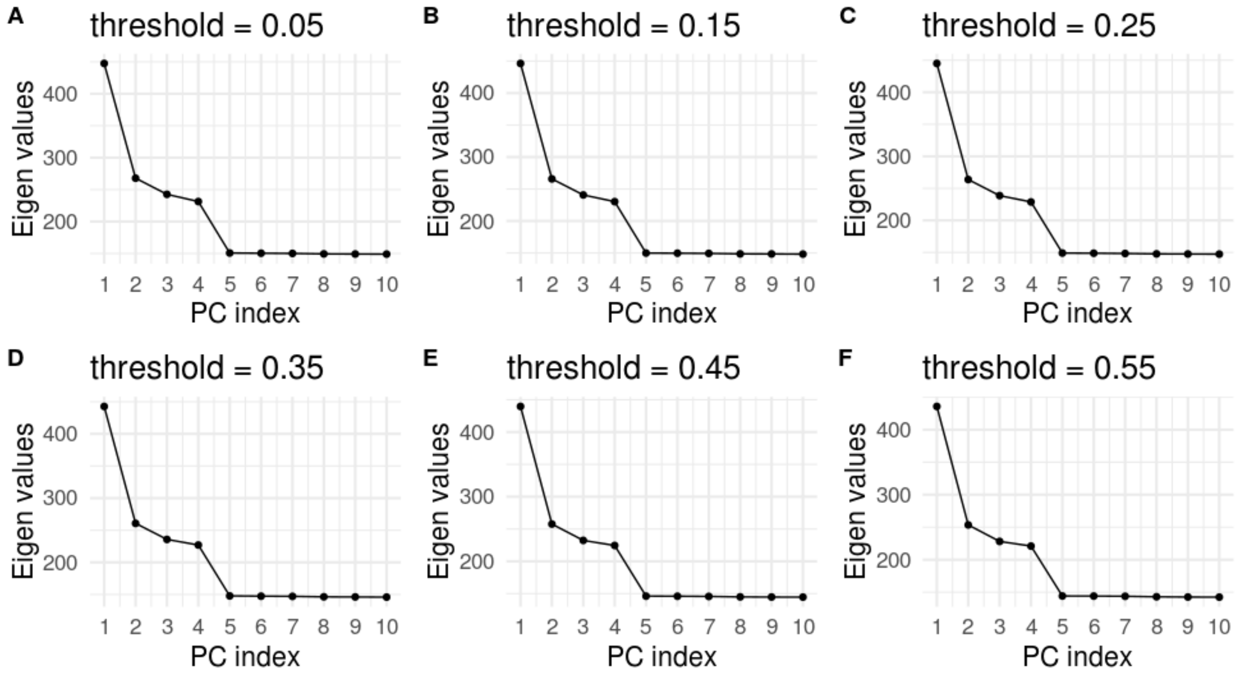


Figure S8: Determining K is robust to variations in accounting for confounders

Scree plot of PCA applied on a D matrix of parameters (n = 100, α0 = 1, $\varepsilon$ = 0.2, G = 1, K = 5) after the removal of confounding probes by linear regression, using different adjusted p-values thresholds. (**A**) p-value < 0.05 (22,758 remaining probes). (**B)** p-value <0.15 (22,532 remaining probes). (**C**) p-value < 0.25 (22,279 remaining probes). (**D**) p-value < 0.35 (21,859 remaining probes). (**E**) p-value < 0.45 (21,400 remaining probes). (**F**) p-value < 0.55 (20,770 remaining probes).

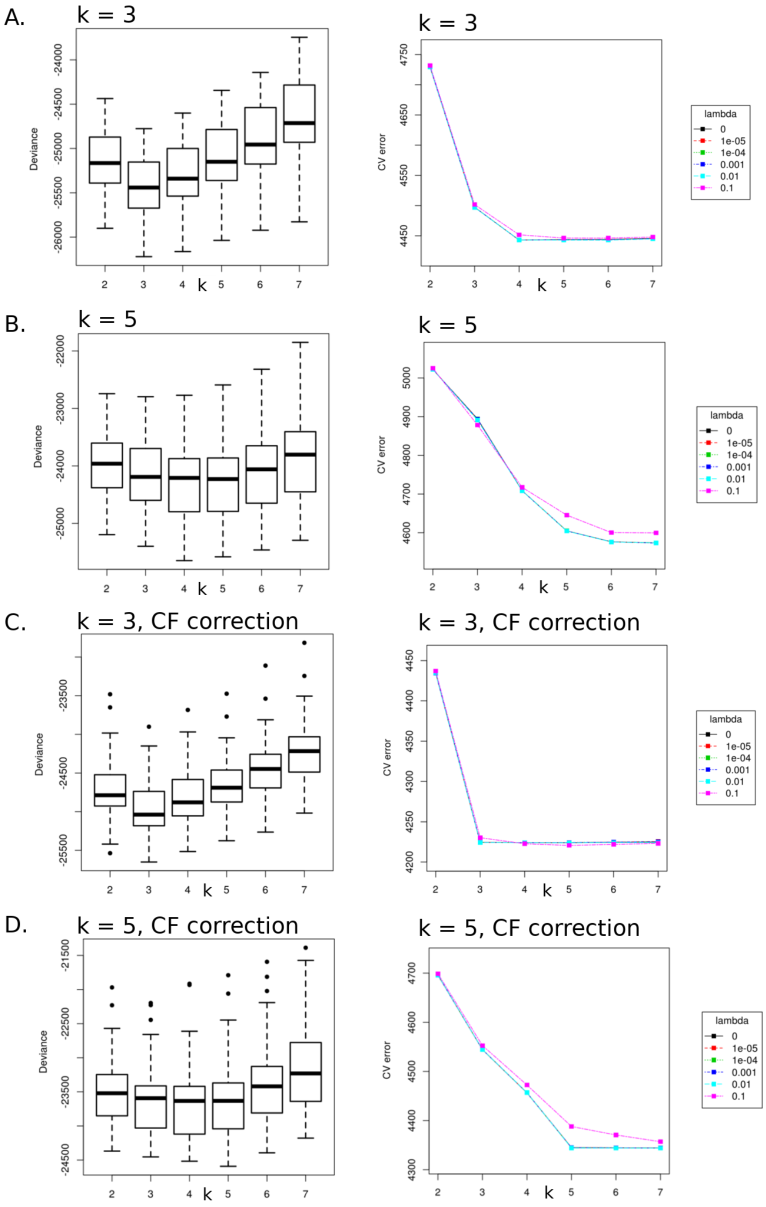


Figure S9 : Determining K by RefFreeEWAS and MeDeCom

To choose K, we also test the bootstrap method of RefFreeEWAS (on the left) and the cross-validation method of MeDeCom (on the right). The D matrix was simulated with the following parameters: n = 100, $\alpha 0 = 1$, $\varepsilon = 0.2$, G = 1 and K = 3 (**A** and **C**) and n = 100, $\alpha 0 = 1$, $\varepsilon = 0.2$, G = 1 and K = 5 (**B** and **D**). (**A**,**B**) Methods applied on **D** matrix before removal of confounding probes (23,381 probes). (**C**, **D**) Methods applied on **D** matrix before removal of confounding probes (22,551 probes in **C**, 22,532 probes in **D**).

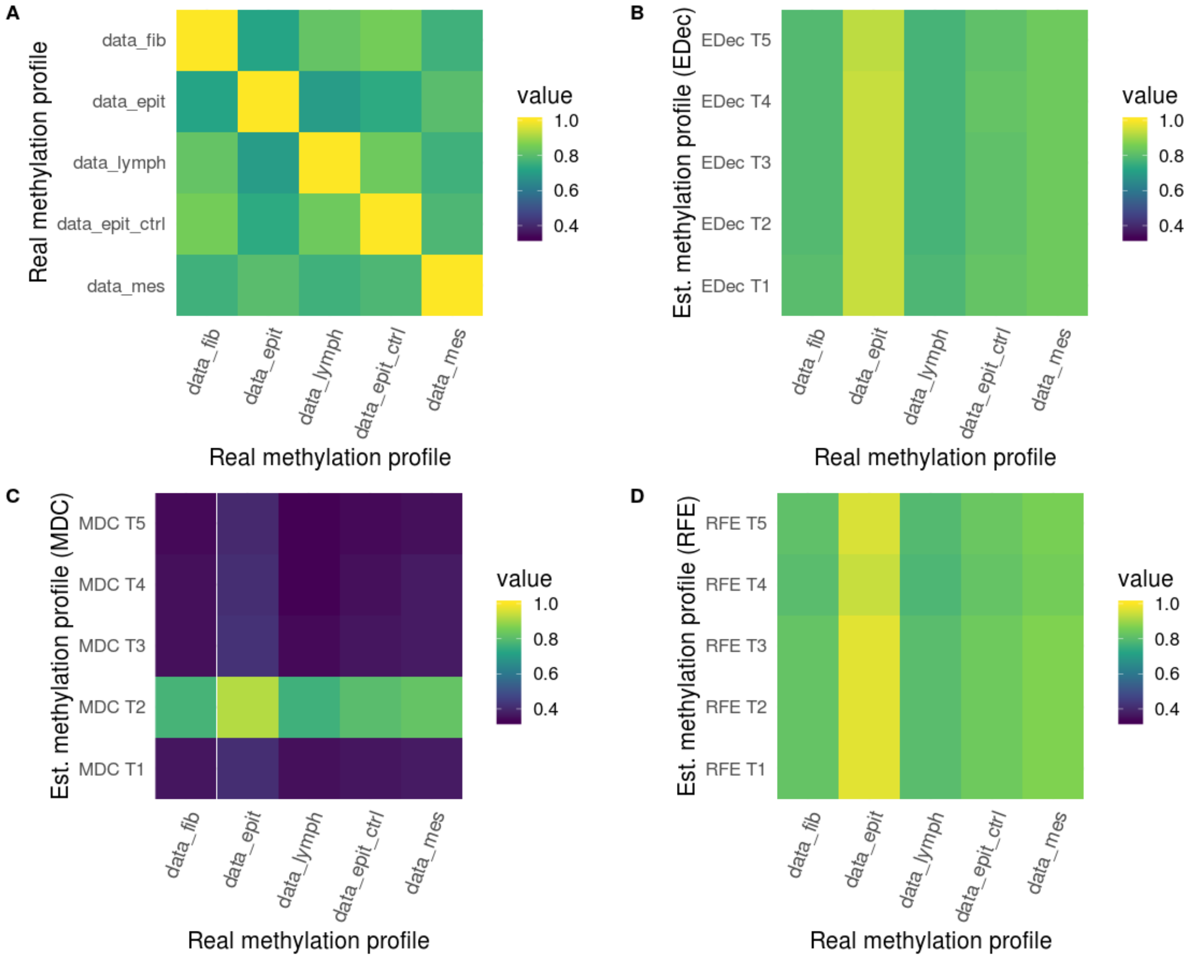


Figure S10: Correlation between estimated and real cell type-specific methylation profiles for a Dirichlet of α0 = 10,000

Heatmap of the correlation between cell type-specific methylation profile used for the simulation and cell type-specific methylation profiles estimated (Est.) by different methods. In (**A**), the correlation between different cell types used for the simulation of the T matrix (data_fib = fibroblast, data_epith = cancerous epithelial, data_lymph = T lymphocytes, data_epit_ctrl = healthy epithelial and data_mes = cancerous mesenchymal). We applied EDec (**B**) , MeDeCom (**C**) and RefFreeEwas (**D**) on a representative simulation of 100 patients (α0 = 10,000, $\varepsilon$ = 0.2, G = 1, K = 5) after the removal of confounding probes by linear regression (22,376 remaining probes). We used the Pearson method to compute the correlation between the estimated cell type-specific methylation profiles and real cell type-specific methylation profiles used for the simulation.

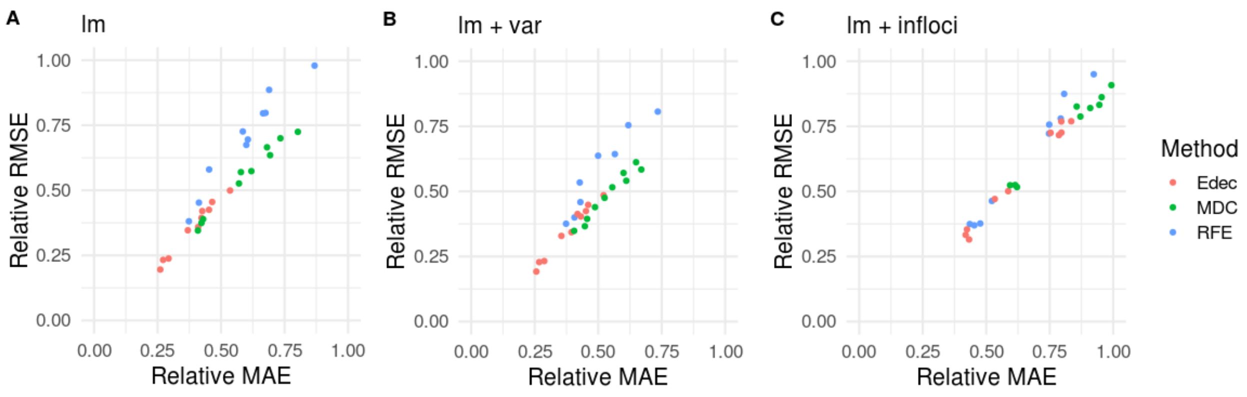


Figure S11: Variation of the error metric: Mean Absolute Error and Root-Mean-Square Error

The Mean Absolute Error (MAE) and the Root-Mean-Square Error (RMSE) were calculated for 10 D matrices (mean value of 10 random noises applied on D) computed from 10 random A. For each pre-treatment method, we compute the relative MAE and the relative RMSE by dividing the error score by that of the matrix before pre-treatment. Each colour represents a different method. RFE stands for RefFreeEWAS, MDC for MeDeCom and EDec for EDec stage 1. T

The following parameters were used to simulate D : K = 5, α0 = 1, $\varepsilon$ = 0.2, G = 1. The methods were run after removal of confounding probes by linear regression (lm, in **A**), after removal of confounding probes and filtering for the most variable probes (lm + var, in **B**), and after removal of confounding probes and filtering of probes expected to biologically vary in methylation levels across constitutive cell types (lm + infloci, in **C**).

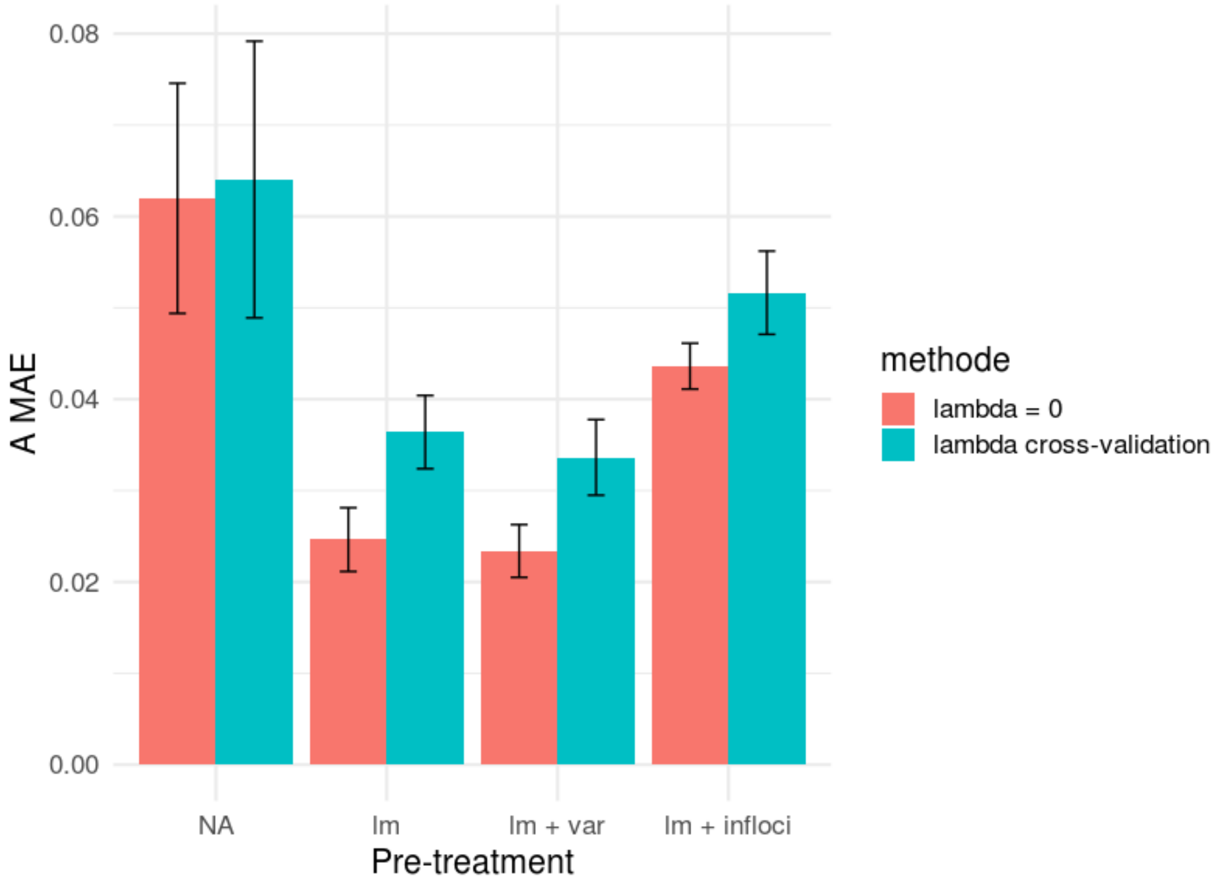


Figure S12: Impact of lambda parameter for MeDeCom

Histogram of `A MAE` (`A MAE`: Mean Absolute Error on estimated A, the matrix of cell proportions) for 10 D matrices (mean value of 10 random noises applied on D) computed from 10 random A. Error bars represent standard deviation on 10 random noises. In red, MeDeCom is run with the parameters lambda = 0. In blue, MeDeCom is run by choosing the best lambda in (0, 0.00001, 0.0001, 0.001, 0.01, 0.1) by cross-validation. The following parameters were used to simulate D : K = 5, α0 = 1, $\varepsilon$ = 0.2, G = 1. The methods were run without pre-treatment (NA), after removal of confounding probes by linear regression (lm), after removal of confounding probes and filtering for the most variable probes (lm + var), and after removal of confounding probes and filtering of probes expected to biologically vary in methylation levels across constitutive cell types (lm + infloci).
