## supplementary_tables for "Guidelines for cell-type heterogeneity quantification based on a comparative analysis of reference-free DNA methylation deconvolution software"

Supplementary Table 1: Number of remaining probes in figure 2

| | No FS | FS variance > 0.02 | FS pval PCA < 0.1 | infloci |
|---|---|---|---|---|
| **n = 20** | 23.381 | 9.875 | 5.957 | 614 |
| **n = 20 + removing of confounding probes** | 21.487 | 9.153 | 5.621 | 591 |
| **n = 100** | 23.381 | 9.593 | 9.017 | 614 |
| **n = 100 + removing of confounding probes** | 22.483 | 8.959 | 8.634 | 607 |

Supplementary Table 2: Number of remaining probes in figure S3

| | No FS | FS variance > 0.02 | FS pval PCA < 0.1 | infloci |
|---|---|---|---|---|
| **n = 20** | 23.381 | 9.082 | 3.953 | 614 |
| **n = 20 + removing of confounding probes** | 21.549 | 8.508 | 3.477 | 597 |
| **n = 100** | 23.381 | 7.742 | 4.271 | 614 |
| **n = 100 + removing of confounding probes** | 22.376 | 7.132 | 3.717 | 605 |

Supplementary Table 3: Mean execution time in figure 7 (minutes)

| | No pre-treatment | Removing of confounding probes | Removing of confounding probes + FS variance > 0.02 | Removing of confounding probes + FS infloci |
|---|---|---|---|---|
| **Edec** | 5.602 | 24.905 | 9.232 | 0.295 |
| **MeDeCom** | 26.848 | 25.549 | 10.227 | 0.765 |
| **RefFreeEwas** | 0.285 | 0.273 | 0.104 | 0.009 |

Supplementary Table 4: Informative loci

cg18509435
cg03608577
cg20542190
cg12242338
cg07349094
cg18963171
cg19680672
cg16869108
cg05788638
cg10917619
cg13445249
cg03602500
cg02311163
cg03872376
cg15374234
cg18841952
cg07950803
cg24355048
cg24423088
cg00463848
cg19486673
cg13897627
cg26135325
cg26530341
cg00041575
cg12417466
cg07374637
cg05615150
cg06183267
cg14366598
cg14153740
cg25890048
cg00895324
cg25119415
cg17657618
cg24338843
cg01637734
cg23338195
cg20856834
cg07525077
cg27431150
cg24272907
cg03914397
cg04574507
cg13765961
cg07548313
cg16812893
cg05767404
cg22325572
cg26799474
cg12910797
cg03547924
cg21949305
cg15720535

cg21453309
cg15475323
cg03171924
cg06469542
cg25509184
cg09061733
cg10942056
cg06490988
cg21129531
cg20308817
cg26200585
cg07197059
cg10574499
cg13705284
cg24019564
cg04527918
cg27635271
cg09440340
cg25431974
cg22764925
cg17791651
cg10266490
cg13641903
cg10257049
cg15901783
cg15083233
cg15149645
cg17465304
cg05724065
cg26521404
cg22396755
cg10282491
cg05606799
cg05590982
cg10046892
cg26884581
cg02324920
cg08044694
cg02838492
cg11052143
cg10280342
cg12782180
cg08519905
cg21488617
cg24989962
cg17349199
cg26606064
cg14859417
cg12513481
cg23606023
cg14371590
cg02490034
cg20373326
cg19138960

cg09037813
cg01222684
cg08314660
cg07498879
cg22972055
cg25313204
cg20622019
cg06465194
cg23797100
cg18345635
cg12041387
cg00661485
cg14754581
cg08077673
cg19539004
cg21614638
cg23663476
cg19863740
cg25947945
cg18997129
cg27433088
cg00503840
cg25949363
cg05670596
cg27016494
cg16363586
cg20583073
cg00499822
cg19248557
cg24861272
cg20080624
cg05656364
cg20876010
cg09954385
cg04947157
cg15164103
cg14236389
cg00412772
cg07816074
cg07665060
cg14417329
cg07359545
cg03386869
cg24541550
cg24619694
cg20356482
cg08615333
cg13053396
cg06630567
cg14137939
cg14918082
cg22979433
cg06589885
cg02506908

cg06290096
cg24365867
cg14027234
cg02586730
cg06339706
cg26215428
cg14297029
cg15679095
cg27256309
cg24315815
cg04106785
cg25182621
cg10857774
cg13521229
cg11213150
cg07260017
cg10222534
cg12069042
cg19466563
cg03389133
cg19664945
cg10635061
cg19433435
cg05955301
cg18003231
cg11042320
cg10362475
cg26924825
cg07168556
cg26220985
cg24447890
cg22416721
cg04999691
cg20368904
cg26453588
cg03870261
cg24340657
cg07354440
cg13030582
cg23130254
cg21577049
cg23092823
cg21815667
cg04797323
cg10756887
cg19352038
cg25465406
cg21790626
cg13035743
cg19797376
cg26620157
cg21233722
cg26189983
cg19358493

cg06825142
cg00347904
cg22709192
cg07696033
cg07634191
cg13577076
cg11108890
cg26069745
cg22341310
cg07533148
cg23587449
cg27188703
cg27009703
cg26113512
cg03874199
cg21238818
cg12265829
cg24516901
cg21529533
cg14056644
cg16761581
cg20616414
cg22377389
cg09516965
cg15540820
cg01335367
cg16158681
cg18536148
cg21604042
cg26416466
cg01683883
cg05345286
cg23887396
cg00363813
cg03421300
cg00744433
cg04245402
cg08687163
cg21459867
cg25612480
cg22477971
cg03109316
cg05445326
cg21484834
cg07785936
cg01731341
cg22190114
cg24870391
cg10883352
cg25527547
cg03977657
cg21624282
cg01835489
cg24835159

cg20556988
cg08458170
cg24147596
cg27622610
cg01580568
cg18462653
cg18223379
cg24833277
cg03332271
cg07908874
cg08886154
cg04711324
cg03116740
cg18623836
cg20324165
cg02833180
cg14333454
cg09847584
cg11206634
cg24276491
cg03684977
cg09936839
cg10145926
cg21201572
cg07711097
cg01119135
cg19686152
cg15422147
cg27341860
cg19831369
cg14851685
cg20797216
cg14209518
cg00757070
cg14404298
cg01441777
cg17233506
cg08214029
cg02237119
cg06848073
cg24134767
cg16609872
cg07705835
cg13300756
cg04329382
cg20484352
cg19258882
cg08124030
cg25426302
cg23499956
cg19759064
cg17471102
cg21602160
cg23579062

cg03003745
cg21250978
cg09307264
cg13439730
cg26672426
cg22889448
cg22580512
cg05245515
cg10052840
cg03752885
cg22131172
cg24855780
cg24620905
cg25946374
cg27324619
cg09152089
cg13928306
cg12603560
cg19923326
cg18053607
cg08578641
cg26143719
cg24765079
cg17826679
cg02293044
cg21663431
cg14519350
cg16509569
cg07380416
cg09902130
cg26285698
cg10590292
cg05751148
cg11600161
cg18384097
cg17936488
cg24497819
cg07973967
cg20792833
cg23612220
cg26158194
cg19663795
cg14145194
cg24545967
cg18621299
cg20425130
cg00447208
cg21898046
cg17771150
cg23506842
cg15258980
cg25671438
cg19252956
cg13821008

cg12177677
cg16068833
cg26149678
cg15691199
cg24088438
cg04275881
cg11804789
cg13470920
cg01348086
cg16749930
cg13273136
cg16777510
cg25866075
cg20994801
cg08818984
cg15679651
cg11201532
cg19005210
cg06855803
cg05989054
cg22407458
cg18611122
cg17199483
cg07376232
cg26928972
cg09682183
cg12108912
cg24450631
cg17709873
cg12836863
cg06144905
cg08804892
cg23352695
cg20018806
cg00168942
cg13634319
cg11098259
cg20070090
cg11484872
cg24625388
cg05501357
cg26267310
cg15840985
cg18940763
cg27365426
cg04797496
cg27562023
cg07086380
cg03548857
cg10934032
cg15387123
cg11203041
cg14114267
cg05246522

cg10037005
cg18908499
cg04301614
cg08585897
cg14451276
cg25229172
cg17998964
cg03294491
cg01001286
cg23213217
cg17527798
cg05130485
cg15645309
cg15261665
cg11304234
cg04915566
cg07935264
cg01280080
cg14448116
cg04228042
cg14902389
cg08529852
cg11761535
cg12564453
cg11998307
cg13354523
cg00597076
cg18493147
cg06910100
cg04995717
cg15127733
cg22844623
cg18533225
cg05064181
cg10275770
cg19537511
cg20967028
cg21012874
cg11398517
cg12876594
cg21142272
cg19368582
cg26245202
cg24664957
cg10157098
cg10061138
cg02254407
cg19242268
cg05046097
cg01169778
cg09243021
cg08779777
cg25902889
cg03636183

cg05050341
cg23679724
cg19731122
cg18771300
cg15407570
cg13797282
cg03483626
cg11136562
cg11484576
cg10569414
cg14289461
cg12456510
cg04123507
cg16501028
cg06003187
cg00350478
cg09299388
cg07039362
cg07613153
cg13412615
cg24919884
cg15903395
cg18552413
cg18153060
cg23444894
cg11939496
cg17983307
cg25363317
cg20647137
cg14269477
cg12032049
cg06403553
cg00601486
cg12200412
cg01087382
cg16666160
cg11812202
cg11481351
cg04498511
cg16581199
cg01671881
cg22077553
cg23514672
cg25093045
cg11465372
cg02423618
cg17561452
cg16084788
cg04618528
cg24030627
cg07830847
cg10377274
cg25141674
cg20485165

cg11234457
cg00745543
cg23749046
cg22988566
cg25737664
cg15554401
cg05260966
cg22445920
cg01469547
cg26628847
cg11113534
cg02807948
cg06849477
cg24457403
cg24697329
cg08203715
cg19370451
cg15481539
cg07753583
cg13379763
cg02737335
cg20488657
cg11706111
cg07879977
cg20543571
cg23382741
cg23767977
cg27108154
cg07478122
cg24024214
cg21256656
cg26504906
cg20661303
cg07728874
cg03112869
cg09995854
cg07947016
cg00510787
cg23865698
cg19324627
cg01869233
cg23627134
cg09747578
cg18022926
cg04081402
cg07165793
cg24293567
cg08012287
cg00201234
cg25288155
cg20664201
cg06821120
cg01110312
cg17602451

cg15195412
cg18015044
cg06980053
cg26561254
cg13755535
cg26608332
cg19815139
cg01138020
cg12228229
cg25827112
cg24691453
cg02735486
cg05656180
cg11432797
cg20725021
cg13904968
cg05163057
cg17974185
cg04008913
cg24269276
